# Ascidian embryonic cells with properties of neural-crest cells and neuromesodermal progenitors of vertebrates

**DOI:** 10.1101/2023.05.24.542044

**Authors:** Tasuku Ishida, Yutaka Satou

**Affiliations:** Department of Zoology, Graduate School of Science, Kyoto University, Sakyo, Kyoto, 606-8502, Japan

**Author notes:** Correspondence to Yutaka Satou.

## Abstract

Neural-crest cells and neuromesodermal progenitors (NMPs) are multipotent cells that are important for development of vertebrate embryos. In embryos of ascidians, which are the closest invertebrate relatives of vertebrates, several cells located at the border between the neural plate and the epidermal region have neural-crest-like properties; hence, the last common ancestor of ascidians and vertebrates may have had ancestral cells similar to neural-crest cells. However, these ascidian neural-crest-like cells do not produce cells that are commonly of mesodermal origin. Here, we showed that a cell population located in the lateral region of the neural plate has properties resembling those of vertebrate neural-crest cells and NMPs. Among them, cells with *Tbx6-related* expression contribute to muscle near the tip of the tail region, and cells with *Sox1/2/3* expression give rise to the nerve cord. These observations and cross-species transcriptome comparisons indicate that these cells have properties like those of NMPs. Meanwhile, transcription factor genes *Dlx.b*, *Zic-r.b*, and *Snai,* which are reminiscent of a gene circuit in vertebrate neural-crest cells, are involved in activation of *Tbx6-related.b*. Thus, the last common ancestor of ascidians and vertebrates may have had cells with properties of neural-crest cells and NMPs, and such ancestral cells may have produced cells commonly of ectodermal and mesodermal origins.

## Introduction

In most animal embryos, three germ layers are specified in early development, and these are further specified to various cell types. However, neural-crest cells, which are formed in the neural plate border of vertebrate embryos, retain or re-activate their ability to produce cells that are commonly of mesodermal and ectodermal origins, even after gastrulation^1–3^. It is widely believed that neural-crest cells contributed to evolution of vertebrates, especially evolution of the head region^4^. Neural-crest cells give rise to various cells including the pharyngeal skeleton, smooth muscle of the aortic arches, and the peripheral neurons. In other words, this cell population produces cells commonly of ectodermal origin and mesodermal origin. The gene regulatory network for differentiation of this cell population has been extensively studied, and its hierarchical structure has been uncovered^5^. Specifically, a gene regulatory circuit typically containing *Dlx5/6*, *Msx1*, *Zic1*, *Tfap2*, and *Pax3/7* acts at the early neural plate border, and soon after it activates genes including *Snai* and *Id* in premigratory neural-crest cells, although there may be small differences among species.

Recent studies have suggested that embryos of ascidians, which belong to the subphylum Tunicata, the sister group of vertebrates, contain cells that share an evolutionary origin with vertebrate neural-crest cells^6–9^. These cells indeed differentiate into cells that include sensory neurons and pigment cells. Although homology of ascidian cells producing sensory bipolar tail neurons and vertebrate neural-crest cells is controversial^10^, it is likely that the last common ancestor of vertebrates and ascidians had cells like vertebrate neural-crest cells. Ascidian neural-crest-like cells identified so far do not produce cell types that are commonly of mesodermal origin, which raises the question of whether ancestral neural-crest cells of the last common ancestor of vertebrates and ascidians (ancestral Olfactores) had the ability to produce cells that were commonly not only of ectodermal origin, but also of mesodermal origin.

Meanwhile, neuromesodermal progenitors (NMPs) of vertebrate embryos, which reside in the tailbud, are another cell population that has the ability to produce mesodermal (presomitic mesodermal) and ectodermal (spinal cord) cells of posterior structures in late embryos^11,12^. NMPs express *T* and *Sox2* simultaneously^13,14^. Mesodermal cells differentiating from NMPs express *Tbx6*, which represses *Sox2*, while neural cells differentiating from NMPs maintain *Sox2* expression^13,15–18^. It has not been determined whether ancestral Olfactores had NMP-like cells. If it had, did ancestral NMP-like cells produce cells that are commonly of mesodermal and ectodermal origins and how did these ancestral neural-crest cells and NMPs contribute to the body plan of ancestral Olfactores?

The central nervous system of ascidian larvae is derived from three pairs of cells of 8-cell embryos, and these pairs and their descendants are called a-, b-, and A-line cells^19–21^. In other words, these three lineages of cells make up the neural plate. However, the most anterior portion of the neural plate is now considered to be the structure that shares an evolutionary origin with vertebrate cranial placodes^22–27^; therefore it may be better to call this region the anterior neural plate border. Likewise, b-line cells constitute the lateral part of the neural plate, and mainly contribute to ependymal cells of the dorsal row of the nerve cord, but not to neurons^19,21^. These cells are derived from a pair of cells called b6.5 at the 32-cell stage. At the 64-cell stage, its daughter cells, b7.9 and b7.10, laterally abut cells that contribute to neurons of the central nervous system (Figure S1). Among their daughter cells, b8.17 and b8.19 (and their descendants) also abut cells that contribute to neurons of the central nervous system, while the other daughter cells (b8.18 and b8.20) and their descendants are located more laterally at the gastrula stage (Figure 1a). The former cells (b8.17 and b8.19) have been regarded as neural plate cells^21^, and we call them and their descendants lateral neural plate cells (LNPCs) in the present study. Peripheral sensory neurons are derived from the latter cells (b8.18/20-line cells), and these cells may share their evolutionary origin with vertebrate neural-crest cells (cyan cells in Figure 1a)^7,8^. In addition, another pair of neural-crest-like cells, which produce pigment cells, is identified in the neural plate (asterisks in Figure 1a)^6^. We investigated the b6.5-descendants in embryos of ascidians (*Ciona robusta*, or *Ciona intestinalis* type A) to answer the question of the evolutionary origin of multipotent neural-crest cells and NMPs, and found that cells of the LNPC lineage have properties of neural-crest cells and NMPs of vertebrates.

**Figure 1.**
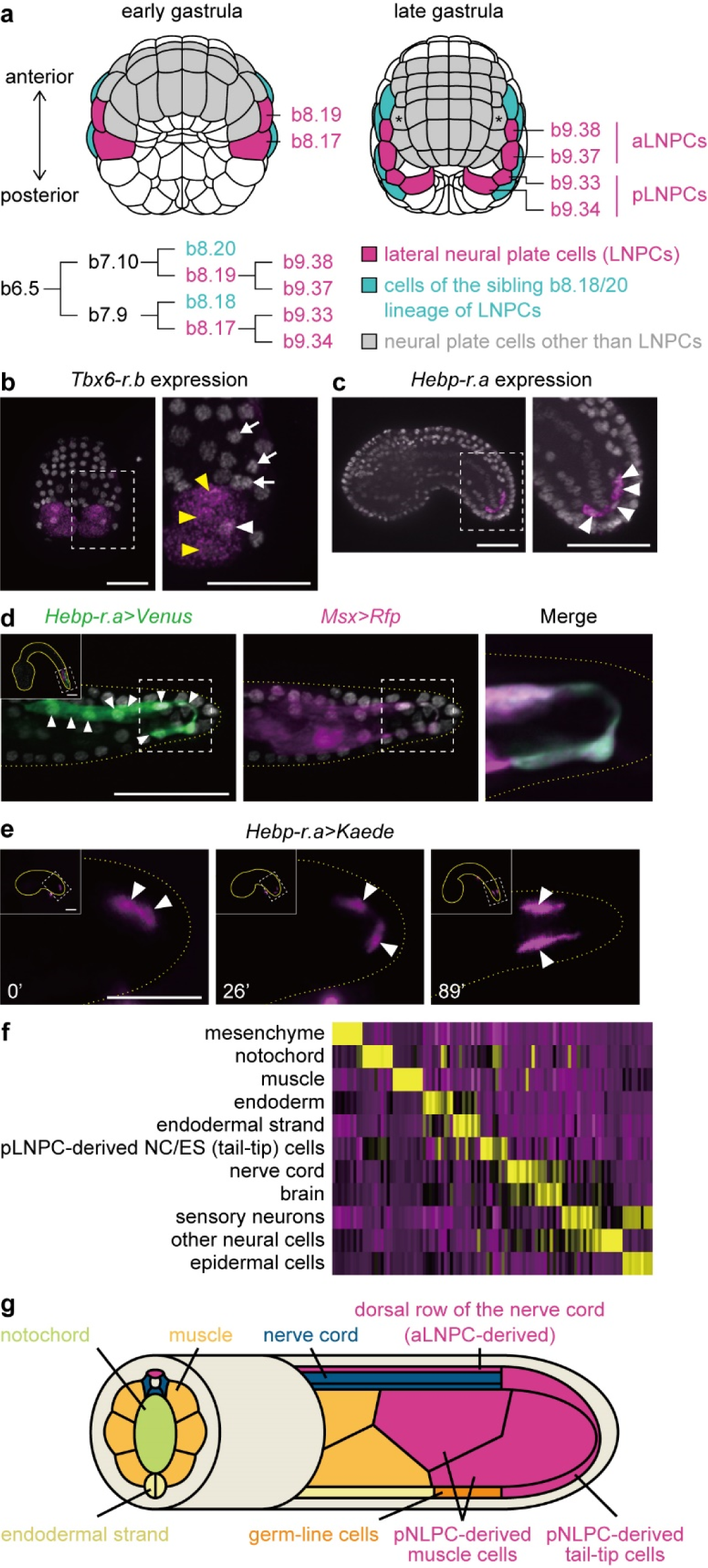
The cell lineage of b8.17 and b8.19 cells. (a) Ascidian bilaterally symmetrical embryos at the early and late gastrula stages. Neural plate cells other than b-line lineage cells are colored gray. Magenta cells are b8.17 and b8.19 at the early gastrula stage, and their descendants at the late gastrula stage, which we call lateral neural plate cells (LNPCs). Cyan cells are b8.18 and b8.20, which are siblings of b8.17 and b8.19, at the early gastrula stage, and their descendants at the late gastrula stage. These cells produce peripheral neurons and are thought to be homologous to vertebrate neural-crest cells^7,8^. In addition, pigment cells in the brain vesicle are derived from cells marked by asterisks, and these cells are also thought to share their origin with vertebrate neural-crest cells^6^. The lineage of LNPCs is shown below. These cells are derived from b6.5 cells in 32-cell embryos. Note that the most anterior part of the neural plate is thought to share its evolutionary origin with vertebrate anterior placodes^22–27^. (b) *Tbx6-r.b* expression revealed by *in situ* hybridization in a middle gastrula embryo (magenta). A higher magnification view is shown on the right. *Tbx6-r.b* expression is observed in b9.34 (white arrowhead). Muscle cells are formed from other lineages and some of these cells also express *Tbx6-r.b* at this stage (yellow arrowheads). LNPCs other than b9.34, which do not express *Tbx6-r.b*, are indicated by white arrows. (c) Expression of *Hebp-r.a* in a tailbud embryo (early tailbud I). A higher magnification view is shown on the right. Cells with *Hebp-r.a* signals are shown by magenta (arrowheads). (d) Expression of reporter constructs that contain upstream regions of *Hebp-r.a* (green) and *Msx* (magenta) in the tip of the tail of a late tailbud embryo (late tailbud II) was examined using specific antibodies against Venus and RFP. Overlaid photographs of signals (left) of the *Hebp-r.a* reporter and DAPI, and (middle) of the *Msx* reporter and DAPI. A higher magnification view of the region near the tip of the tail are shown on the right as an overlaid photograph of the *Hebp-r.a* reporter and the *Msx* reporter. Nuclei of eight cells labelled with *Hebp-r.a* reporter expression are shown by arrowheads in the left photograph. Dorsal is up, and ventral is down. (e) A tailbud embryo expressing *Hebp-r.a>Kaede* reporter was subjected to UV-irradiation for photoconversion of Kaede fluorescence at the early tailbud stage (0 min; early tailbud I). One cell with photoconverted Kaede changed its location to the posterior end at the middle tailbud stage (26 min; middle tailbud I), and to the ventral side at the late tailbud stage (89 min; late tailbud I). Note that only two cells are labelled because of mosaic incorporation of the reporter construct. Photographs at other time points are shown in Figure S3. (f) A heatmap showing gene expression of various cell types. Cell groups labelled ‘nerve cord’ and ‘endodermal strand’ do not include pLNPC-derived nerve cord (NC) or endodermal strand (ES) cells. Note that we do not discriminate four rows of the nerve cord in this analysis. Each vertical line represents expression levels of a single gene. Expression levels were calculated as averages of expression levels of cells in each cell type. The top 10 genes for each cell type are shown. Yellow and magenta represent high and low expression levels, respectively. Photographs in b, c, d, and e are z-projected image stacks overlaid in pseudocolor. In b, c, and d, nuclei are stained with DAPI (gray). Brightness and contrast of photographs in d and e were linearly adjusted. Scale bars, 50 μm. (g) Depiction of the ascidian tail. LNPC-derived cells are colored magenta.

## Results

### Posterior LNPCs give rise to muscle cells and cells near the tip of the tail region

Previous studies^19,28^ have shown that b8.17 (posterior LNPCs, pLNPCs) give rise to cells near the tip of the tail of tailbud embryos. These cells are two pairs of muscle cells and four cells of posterior parts of the nerve cord and endodermal strand. Because *Tbx6-related.b* (*Tbx6-r.b*) is required for specification of all muscle cells, including muscle cells derived from b8.17^29^, we examined *Tbx6-r.b* expression in LNPCs, and found that the posterior daughter cells (b9.34) of b8.17 express *Tbx6-r.b* (Figure 1b). Indeed, these cells expressed *Acta.a*, which encoded muscle actin (Figure S2). Therefore, it is highly likely that the posterior daughters give rise to muscle cells.

This observation implies that pLNPC-derived nerve-cord and endodermal-strand cells are derived from the anterior daughter of b8.17 (b9.33). *Hand-r* is expressed in cells near the tip of the tail^30–33^, but it cannot be used as a specific marker, because it is also expressed in other cells. Therefore, we used *Hand-r* as a marker gene to find genes expressed specifically in pLNPC-derived nerve-cord and endodermal-strand cells in single-cell RNA-sequencing (scRNA-seq) data published previously^34^ (Table S1). Among these genes, we confirmed that *Hebp-r.a* is indeed expressed in putative pLNPC-derived nerve-cord and endodermal-strand cells by *in situ* hybridization (Figure 1c). When we introduced a reporter gene containing the upstream regulatory region of *Hebp-r.a* with another reporter gene containing the upstream region of *Msx*, these two reporters were expressed in the same cells at the tip of the tail region, which is commonly called the tailbud (Figure 1d). Because *Msx* is specifically expressed in b8.17, b8.18, b8.19, and b8.20 in early gastrulae^8,30,35,36^, and because only b8.17 contributes to pLNPC-derived nerve-cord cells and endodermal-strand cells^19^, we conclude that *Hebp-r.a* specifically marks pLNPC-derived nerve-cord cells and endodermal-strand cells. Four cells expressed *Hebp-r.a* at the early tailbud stage (Figure 1c). This coincides with the number of b8.17 derivatives in a previous study^19^. It is likely that these pLNPC-derived nerve-cord and endodermal-strand cells divided once between the early and late tailbud stages, because eight cells expressed *Hebp-r.a* reporter at the late tailbud stage (Figure 1d). Thus, the anterior daughter (b9.33) of pLNPC (b8.17) gives rise to the most posterior parts of nerve-cord cells and endodermal-strand cells and the posterior daughter (b9.34) gives rise to muscle cells.

At the early tailbud stage, pLNPC-derived nerve-cord cells and endodermal-strand cells were initially located in the posterior part of the nerve cord (Figure 1e). Subsequently, some of them changed their location to the ventral side, and these cells were finally located in the posterior part of the endodermal strand (Figure 1e; Figure S3). Using scRNA-seq data^34^, we compared expression profiles of pLNPC-derived nerve-cord cells and endodermal-strand cells with those of other cell types during the larval stage, and found that the transcriptome of pLNPC-derived nerve-cord cells and endodermal-strand cells was quite different from those of the other endodermal strand and nerve cord cells and of other tissues (Figure 1f). This was further supported by the observation that pLNPC-derived nerve-cord cells and endodermal-strand cells did not express a pan-neural marker, *Celf3.a*^37^, or an endodermal-strand marker, *Slc39a-related*^32^ (Figure S4). Accordingly, these pLNPC-derived cells constitute a cell population distinct from nerve-cord cells or endodermal-strand cells located anteriorly.

The ascidian larval nerve cord mostly consists of four rows of cells, while the most posterior part of the nerve cord consists of a row of several cells, but not of four cell rows^38^. Similarly, the endodermal strand mostly consists of two rows of cells, while the most posterior part of the endodermal strand does not, and germ-line cells intervene between the anterior two rows of endodermal strand cells and the most posterior part of the endodermal strand^38^. Judging from their locations, it is likely that these posterior nerve-cord cells and endodermal-strand cells are pLNPC-derived cells. These observations also suggest that these cells may constitute a cell population distinct from nerve-cord cells or endodermal-strand cells located anteriorly (Figure 1g). We call these cells tail-tip cells hereafter, although we do not necessarily rule out the possibility that these cells are specialized cells of the nerve cord or endodermal strand.

### *Tbx6-r.b* is activated by a set of regulatory factors similar to a set that specifies the neural plate border and neural-crest cells in vertebrate embryos

Next, we examined how *Tbx6-r.b* is activated in b9.34. Previous studies have shown that orthologs for genes specifying neural plate border cells and neural-crest cells in vertebrate embryos are expressed in cells including LNPCs of ascidian embryos^30,36,39–41^. We confirmed that *Dlx.b, Msx*, *Snai*, *Zic-r.b*, *Tfap2-r.b*, *Pax3/7*, *Ets1/2.b*, *Lmx1*, and *Id.b* were indeed expressed in LNPCs (Figure S5).

*Dlx.b* is expressed in animal hemisphere cells, including LNPCs, in gastrulae, and is important for fate decision of these cells^39^. *Msx* is expressed in LNPCs and their sibling b8.18/20-line cells^30^, and is required for differentiation of sensory neurons^8,35^. While knockdown of *Msx* did not affect *Tbx6-r.b* expression, *Dlx.b* knockdown downregulated *Tbx6-r.b* in b9.34 (Figure 2a). Note that signals for *Tbx6-r.b* expression in A-line muscle cells are seen over b9.33 cells where *Tbx6-r.b* is not expressed (Figure S6). *Snai* was also expressed in LNPCs (Figure S5), and knockdown of *Snai* downregulated *Tbx6-r.b* expression (Figure 2a). In addition, *Zic-r.b* was downregulated in *Dlx.b* morphants (Figure 2b), and *Snai* was down-regulated in *Zic-r.b* morphants (Figure 2c). Thus, orthologs of genes that specify fates of the neural plate border and neural-crest cells in vertebrate embryos, also regulated *Tbx6-r.b* expression in b9.34.

**Figure 2.**
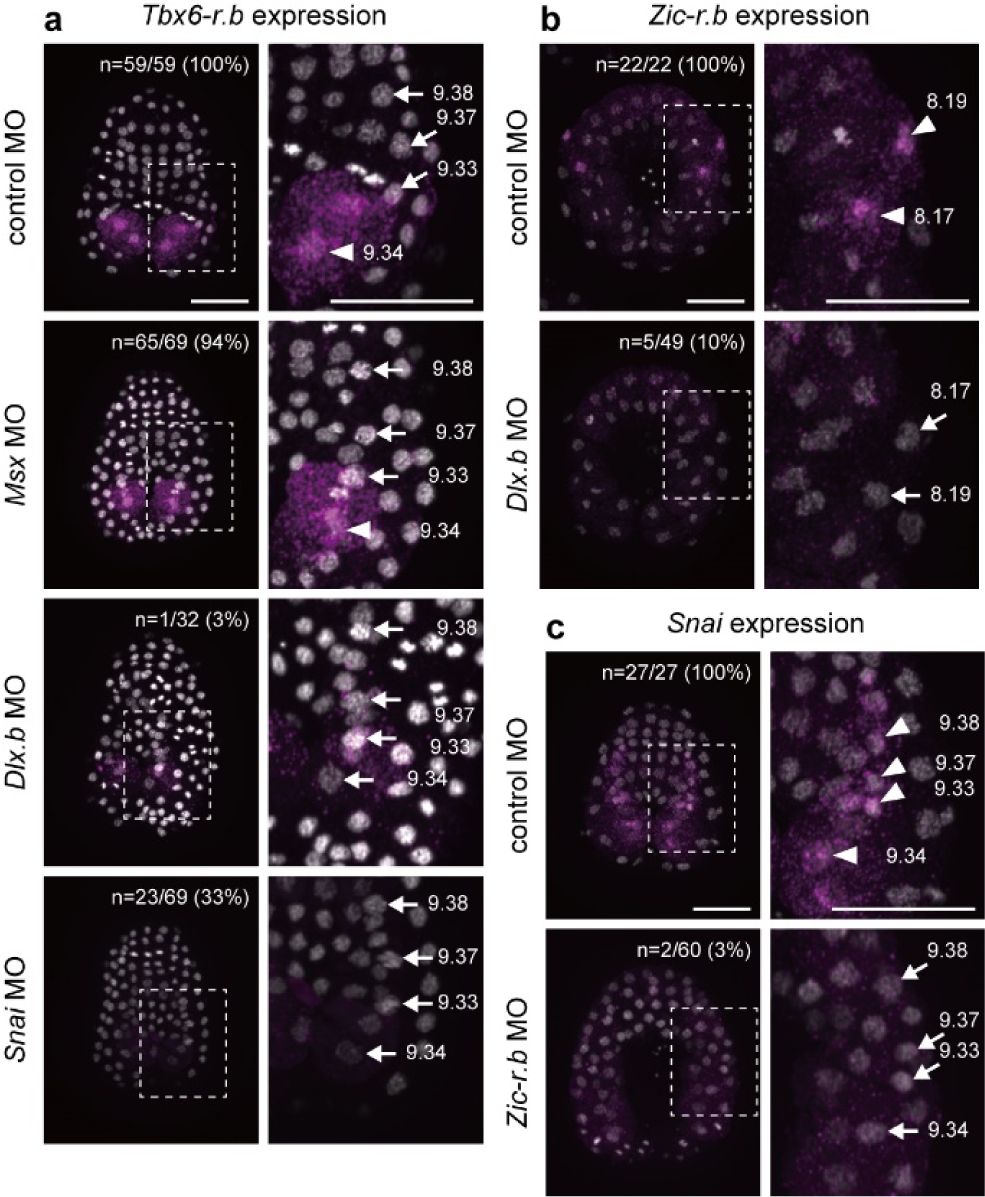
The regulatory gene circuit that activates *Tbx6-r.b* in b9.34. (a) Expression of *Tbx6-r.b* in a control embryo and embryos injected with an MO against *Msx, Dlx.b,* or *Snai*, at the early neurula stage. (b) Expression of *Zic-r.b* in a control embryo and an embryo injected with the *Dlx.b* MO at the early gastrula stage. (c) Expression of *Snai* in a control and an embryo injected with the *Zic-r.b* MO at the middle gastrula stage. Higher magnification views are shown on the right. LNPCs that express and do not express designated genes are indicated by arrowheads and arrows, respectively. Nuclei are stained with DAPI and are shown in gray. Photographs are z-projected image stacks overlaid in pseudocolor. Brightness and contrast of photographs in embryos showing *Tbx6-r.b* expression, except the *Snai* morphant embryo, were adjusted linearly. Numbers of embryos examined and embryos that expressed (a) *Tbx6-r.b* in b9.34, (b) *Zic-r.b* in b8.17 and b8.19, and (c) *Snai* in LNPCs are shown in the panels. In the *Msx* and *Dlx.b* morphants shown in (a), signals for *Tbx6-r.b* expression in A-line muscle cells are seen over b9.33 cells, in which *Tbx6-r.b* is not expressed (see Figure S6). Scale bars, 50 μm.

### *Tbx6-r.b* potentially suppresses *Sox1/2/3* expression

While LNPCs are located in the lateral region of the neural plate and abut epidermal territory near the dorsal side of the blastopore in early gastrulae (see Figure 1a), pLNPC-derived cells are found near the tip of the tail region in later embryos (see Figure 1g). pLNPCs also contribute to muscle cells, in which *Tbx6-r.b* is expressed (see Figure 1b), and *Tbx6-r.a*, a paralog of *Tbx6-r.b*, is expressed in b9.33 and b9.34 (Figure S7a). In addition, anterior LNPCs (aLNPCs; b9.37 and b9.38) contribute to the dorsal row of the nerve cord (see Figure 1g)^19^. These features of LNPCs are evocative of vertebrate NMPs.

Tbx6 is expressed in NMP-derived mesodermal cells of vertebrate embryos and represses *Sox2*, which is highly expressed in NMP-derived neural cells^15,16,18^. In ascidian early embryos, *Sox1/2/3* is expressed in animal hemisphere cells including b6.5, which gives rise to LNPCs^30,39,42^. Indeed, *Sox1/2/3* was expressed in pLNPCs of middle gastrula embryos (Figure S8a), in which *Tbx6-r.a* and *Tbx6-r.b* are expressed (see Figure 1b and Figure S7). However, this *Sox1/2/3* expression became weak at the early neurula stage (Figure S8b). Therefore, we used a probe designed to hybridize with the first intron of *Sox1/2/3* to detect *Sox1/2/3* transcription. In normal unperturbed embryos, while nascent *Sox1/2/3* RNA was detected only in aLNPCs and in cells that contribute to the brain, it was not detected in pLNPCs (Figure S8c). When we overexpressed *Kaede* using the upstream regulatory region of *Msx* in LNPCs and their sibling b8.18/20-line cells (*Msx>Kaede*) as a control, expression of nascent *Sox1/2/3* transcripts was not changed (Figure 3a). On the other hand, when we overexpressed *Tbx6-r.b* using the upstream regulatory region of *Msx* (*Msx>Tbx6-r.b*), nascent *Sox1/2/3* transcripts were not detected in LNPCs, whereas expression in cells that contribute to the brain, in which *Tbx6-r.b* was not overexpressed, was unaffected (Figure 3a). *Lmx1*, which is normally expressed in aLNPCs of neurula embryos under control of *Msx*, *Nodal* and *Zic-r.b*, and contributes to specification of the dorsal row of the nerve cord by activating *Pax3/7*^36,43^, was also downregulated (Figure 3b). Instead, these cells ectopically expressed a muscle marker gene, *Acta.a* (Figure 3c). Thus, aLNPCs have the potential to become muscle cells, and a key determinant for muscle fate, *Tbx6-r.b*, potentially represses *Sox1/2/3*.

**Figure 3.**
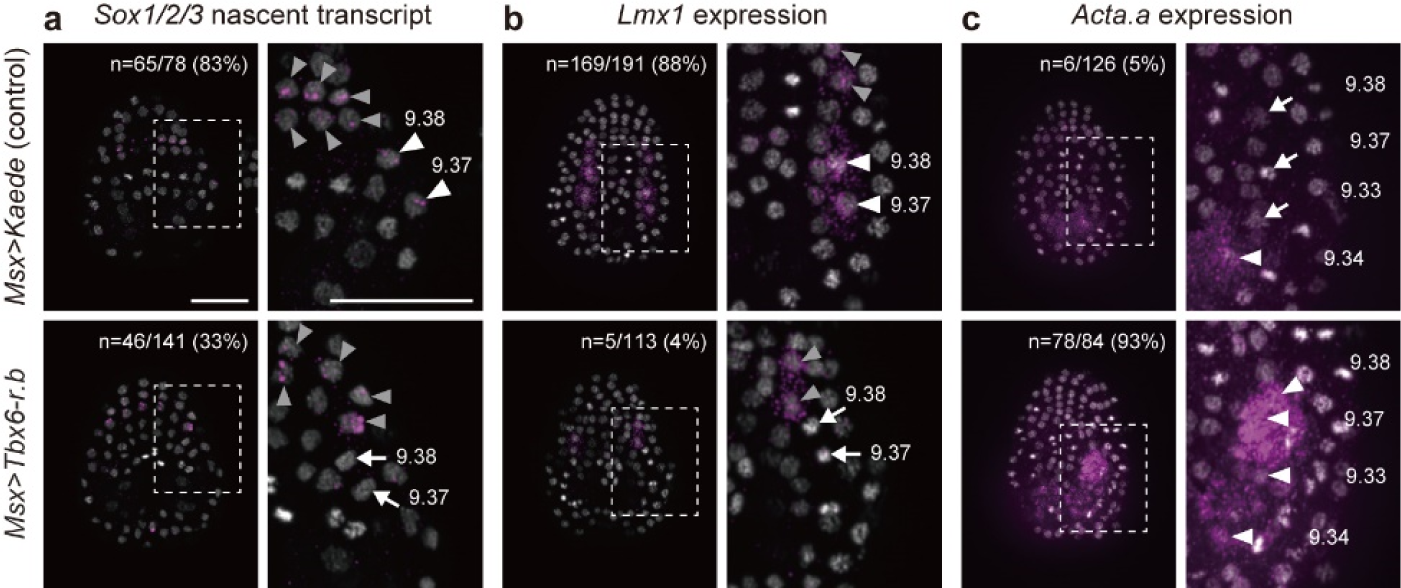
*Tbx6-r.b* can repress *Sox1/2/3* in aLNPCs and can change their fate to muscle. (a) *Sox1/2/3* is transcribed in aLNPCs (b9.37 and b9.38; white arrowheads) and in cells that contribute to the central nervous system (gray arrowheads) of control middle gastrula embryos introduced with *Msx>Kaede*. On the other hand, *Sox1/2/3* is not transcribed in aLNPCs in embryos introduced with *Msx>Tbx6-r.b* (white arrows), while transcription in cells that contribute to the central nervous system is unaffected (gray arrowheads). A probe designed to hybridize with the first intron of *Sox1/2/3* was used to detect *Sox1/2/3* nascent transcripts, and signals are typically seen as one or two spots in nuclei. (b) While *Lmx1* is expressed in aLNPCs (white arrowheads) of control early neurula embryos expressing *Msx>Kaede*, it is not expressed in these cells of embryos expressing *Msx>Tbx6-r.b* (white arrows). Note that *Lmx1* expression in cells that contribute to the central nervous system is unaffected (gray arrowheads). (c) In control early neurula embryos introduced with *Msx>Kaede*, *Acta.a*, which encodes a muscle actin, is expressed in b9.34, while it is not expressed in the remaining LNPCs. In embryos introduced with *Msx>Tbx6-r.b*, *Acta.a* is expressed in all LNPCs. Arrowheads and arrows indicate LNPCs with and without *Acta.a* expression, respectively. Photographs are z-projected image stacks overlaid in pseudocolor; magenta, *in situ* hybridization signals; gray, nuclei stained with DAPI. Brightness and contrast of photographs were adjusted linearly. Numbers of embryos examined and embryos that expressed designated genes (a, b) in LNPCs and (c) in b9.33/37/38 are shown in the panels. Scale bars, 50 μm.

We also examined *Sox1/2/3* expression in morphants of *Msx*, *Snai*, *Dlx.b*, or *Zic-r.b*, and found that *Dlx.b* and *Zic-r.b*, but not *Msx* or *Snai*, are required for *Sox1/2/3* transcription in aLNPCs (Figure S9).

### Single-cell transcriptome data support similarity of ascidian LNPCs and vertebrate NMPs

The location of LNPCs in late embryos and the regulatory circuit of *Tbx6-r.b* and *Sox1/2/3* support the hypothesis that LNPCs in ascidian embryos share an evolutionary origin with vertebrate NMPs. To further confirm this hypothesis, we compared transcriptomes of LNPCs with those of zebrafish embryonic cells using scRNA-seq data published previously^34,44^. To perform a cross-species comparison, we identified ortholog groups using OrthoFinder^45^ between genes of *Ciona* and zebrafish, and compared ascidian LNPCs with all available cells of zebrafish embryos from 10 to 24 hpf. Ascidian notochord cells and muscle cells were also included as controls. We performed clustering of cells at three different resolutions (Table S2-S5). At the highest resolution, all *Ciona* notochord cells and 56% of zebrafish notochord cells were found together in a single cluster, and all *Ciona* muscle cells and 97% of zebrafish myotome and muscle cells were found together in three clusters (Figure S10). This observation indicates that our cross-species comparison successfully grouped cells according to cell types.

Similarly, at the middle resolution, all *Ciona* notochord cells and 83% of zebrafish notochord cells were found together in a single cluster, and all *Ciona* muscle cells and 99% of zebrafish myotome and muscle cells were found together in a single cluster (Figure 4ab; Figure S11; Figure S12). At this resolution, pLNPCs (b9.33 and b9.34) and tail-tip cells (b9.33 derivatives) were found in cluster 13 (Figure 4ace). This cluster contains presomitic-mesoderm cells in the tailbud of zebrafish embryos (Figure 4ade). Zebrafish tailbud presomitic-mesoderm cells were included not only in cluster 13 but also in cluster 9 and cluster 12 (Figure 4e). Cluster 12 mainly consists of myotome and muscle cells. Because almost all cells (99.8%) in clusters 9 and 13 constitute a single cluster at a low resolution (Figure S13; Table S2), clusters 9 and 13 are close to each other. These results indicate a close relationship between ascidian b9.33/b9.34/pLNPC-derived cells and zebrafish tailbud-presomitic-mesoderm cells.

**Figure 4.**
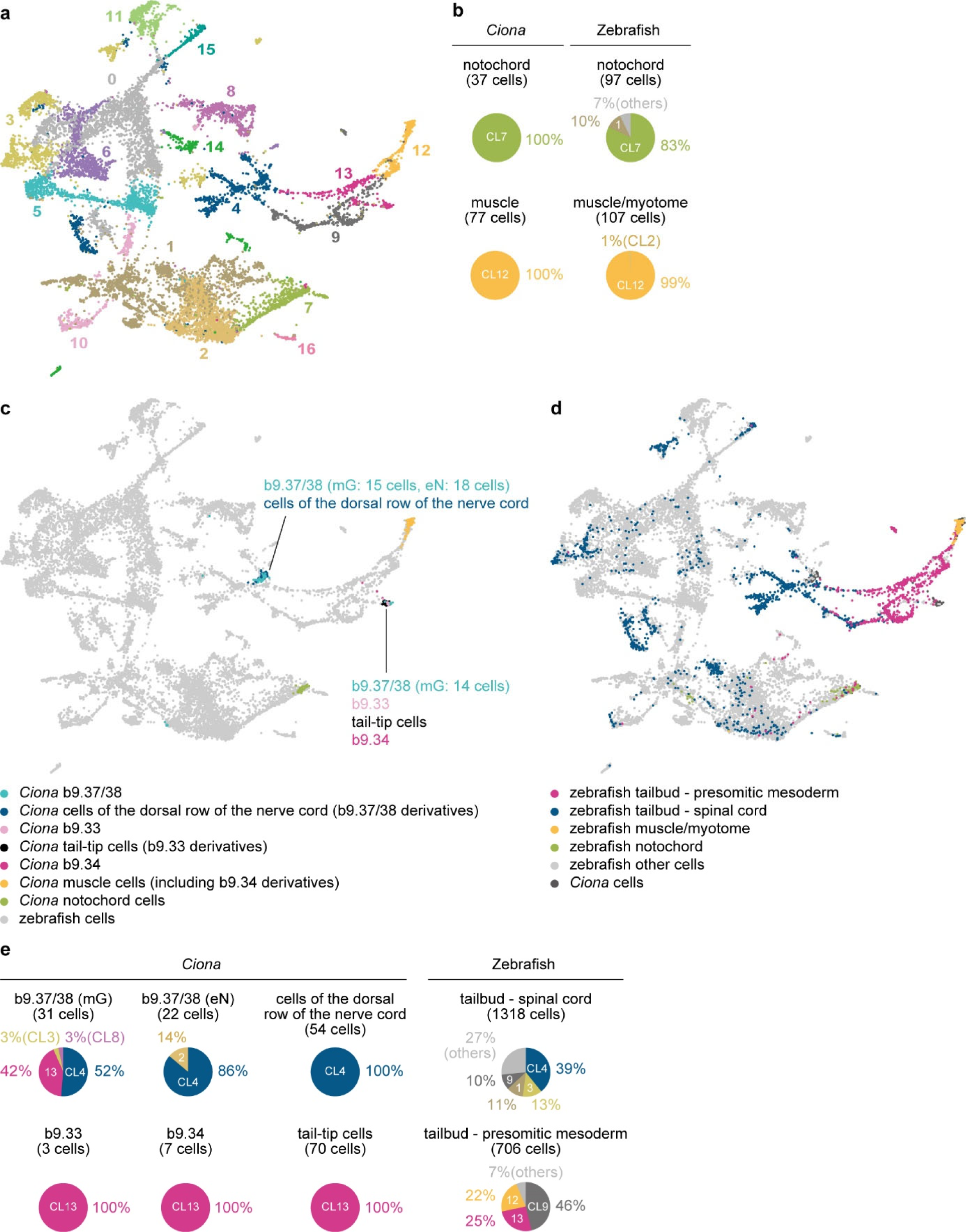
A cross-species comparison of single-cell transcriptome data indicates affinities between ascidian LNPCs and tailbud cells of zebrafish embryos. (a) A UMAP plot that includes single-cell transcriptome data of 10 to 24 hpf embryos of zebrafish, and LNPCs and their derivatives in ascidian embryos from the middle gastrula to larval stages. For controls, muscle cells and notochord cells of ascidian middle tailbud embryos are also included. Different clusters are indicated by different colors. (b) Pie charts show that notochord and muscle cells of *Ciona* and zebrafish are largely grouped into the same clusters. (c) *Ciona* LNPCs, muscle cells, and notochord cells are colored in the same UMAP as that shown in a. (d) Zebrafish tailbud cells, muscle cells, notochord cells are colored in the same UMAP as that shown in a. (e) Proportions of clusters that include designated cells are shown by pie charts.

Nearly half of b9.37 and b9.38 cells of middle gastrula embryos (42%) were found in cluster 13, which included b9.33, b9.34, and tail-tip cells, and almost all of the remaining cells (52%) were found in cluster 4 (Figure 4ace). At the early neurula stage, 86% of of b9.37 and b9.38 cells were found in cluster 4 (Figure 4ace). At later stages, all b9.37/b9.38 derivatives (cells in the dorsal row of the nerve cord) were found in cluster 4 (Figure 4ace). This cluster contains spinal cord cells of the tailbud of zebrafish embryos (Figure 4ade). In fact, among all spinal cord cells of the tailbud of zebrafish embryos, 39% of cells were found in cluster 4 (Figure 4e). This result indicates a close relationship between ascidian b9.37/b9.38-lineage cells and zebrafish tailbud spinal-cord cells. In other words, our analysis using single-cell transcriptome data supports the hypothesis that LNPCs have properties similar to those of zebrafish NMPs, which reside in the tailbud and contribute to the spinal cord and somites.

## Discussion

Our data show the similarity between ascidian LNPCs and vertebrate neural plate border cells that give rise to neural-crest cells. This suggests that ascidian LNPCs share an evolutionary origin with vertebrate neural-crest cells. First, both ascidian and vertebrate cell populations arise in the lateral border (or lateral side) of the neural plate. Second, most genes for transcription factors that specify the neural plate border and neural-crest cells in vertebrates are expressed in these ascidian cells. These genes include *Dlx.b, Msx*, *Snai*, *Zic-r.b*, *Tfap2-r.b*, *Pax3/7*, *Ets1/2.b*, *Lmx1*, and *Id.b*, and we showed that the gene circuit of *Dlx.b*, *Zic-r.b,* and *Snai* indeed regulates *Tbx6-r.b* in pLNPCs. In addition, in aLNPCs, *Msx* and *Zic-r.b* regulate *Lmx1*, and *Lmx1* regulates *Pax3/7*^36^.

LNPCs have been regarded as part of the neural plate^19,21^. However, on the basis of our data, as well as the observation that LNPCs do not contribute to central nervous system neurons^19,21^, we propose to call LNPCs lateral neural plate border cells. Because their sibling b8.18/20 lineage contributes to sensory neurons in the tail and because these cells may share an evolutionary origin with vertebrate neural-crest cells^7,8^, the entire b6.5-lineage could be a cell population that shares an evolutionary origin with neural plate border cells of vertebrates.

On the other hand, our data also showed the similarity between ascidian LNPCs and vertebrate NMPs. *Sox1/2/3* is initially expressed in all animal hemisphere cells, including cells giving rise to LNPCs^30,39,42^, and our data indicate that it is downregulated in pLNPCs. In vertebrate NMPs, cells with strong *Tbx6* expression contribute to mesodermal cells and cells with strong *Sox2* expression contribute to the posterior part of the neural tube^13,16–18^. In ascidian embryos, b9.34 expresses *Tbx6-r.b* and gives rise to muscle cells. Another pLNPC, b9.33, expresses *Tbx6-r.a*, but produces tail-tip cells. On the other hand, aLNPCs continue to express *Sox1/2/3* and give rise to the nerve cord. The ascidian larval tail nerve cord does not contain neurons^46^, and it is debatable which part of the vertebrate central nervous system corresponds to the ascidian nerve cord^23,36,47,48^. Nevertheless, cells near the tip of the tail with *Sox2* (or its ortholog) expression contribute to the posterior part of the central nervous system in both ascidian and vertebrate embryos. In addition, in both embryos, *Tbx6* (or its ortholog) negatively regulates *Sox2* (or its ortholog) and promotes mesodermal fate. These shared properties indicate a common evolutionary origin of vertebrate NMPs and ascidian LNPCs. Consistently, our cross-species comparison of zebrafish and *Ciona* scRNA-seq data showed similarity between ascidian LNPCs and vertebrate NMPs.

While NMPs are characterized by co-expression of *Sox2* and *T* in vertebrate embryos^13^, *T* is not expressed in LNPCs or in any cells with *Sox1/2/3* expression in ascidian embryos^49^. *T* is expressed near the blastopore in many animals, including deuterostomes and protostomes, and is used as a pan-mesodermal marker in vertebrate embryos^50^. In ascidian embryos, this conserved expression is lost and *T* is expressed exclusively in the notochord lineage^49^. Therefore, it is possible that ancestral cells of ascidian LNPCs and vertebrate NMPs expressed *T* and that this expression was lost in the ascidian lineage, although it is equally possible that *T* expression was acquired in the vertebrate lineage. Nevertheless, data in the present study strongly indicate that the evolutionary origin of NMPs dates back to a common ancestor of ascidians and vertebrates (ancestral Olfactores).

Thus, LNPCs have properties of vertebrate neural-crest cells and NMPs. This suggests that the ancestral Olfactores may have had a cell population resembling ascidian LNPCs, and that such ancestral cells may have evolved into neural-crest cells and NMPs in the vertebrate lineage. Meanwhile, because cells of the LNPC-sibling b8.18/20 lineage are thought to be homologs of neural-crest cells^7,8^, it is possible that the ancestral Olfactores also had additional neural-crest-like cells that did not have NMP-like properties.

While ascidian neural-crest-like cells identified in previous studies produce sensory neurons and pigment cells^6–8^, LNPCs give rise not only to cells that are normally of ectodermal origin, but also cells that are normally of mesodermal origin. Contrary to the previous hypothesis^51–54^, this observation indicates that the evolutionary origin of multipotency of vertebrate neural-crest cells may date back to the ancestral Olfactores, although our results are not mutually exclusive to the hypothesis^55^ that various mesodermal developmental programs have been co-opted under gene regulatory networks for neural-crest cells and NMPs in the vertebrate lineage after the spilt of vertebrates and tunicates.

Several studies have shown a close relationship between trunk neural-crest cells and NMPs using human pluripotent cultured cells^56–58^. In addition, trunk neural-crest cells are thought to have emerged earlier than other neural-crest cells^59,60^. Therefore, ancestral cells of neural-crest cells may have had properties of NMPs, and ancient Olfactores may have had such cells. *Ciona* LNPCs, which have properties of neural-crest cells and NMPs, may represent such an ancestral state, and multipotency of these two cell populations may share a common evolutionary origin.

## Acknowledgements

We thank Reiji Masuda, Shinichi Tokuhiro, Chikako Imaizumi (Kyoto University), Manabu Yoshida (University of Tokyo), and other members working under the National BioResource Project for *Ciona* (MEXT, Japan) at Kyoto University and the University of Tokyo for providing experimental animals. This research was supported by grants from the Japan Society for the Promotion of Science under the grant numbers 21H02486 and 21H05239 to YS. The manuscript was edited by a technical editor, Steven D. Aird.

## Author contribution

TI performed experiments. YS drafted the original manuscript. TI and YS reviewed the manuscript draft, revised it, and approved the final version.

## Data availability

All data generated or analyzed during this study are included in this published article.

## Materials and Methods

### Animals and gene identifiers

Adult specimens of *Ciona intestinalis* (type A; also called *Ciona robusta*) were obtained from the National BioResource Project for *Ciona*. cDNA clones were obtained from our EST clone collection^31^. Identifiers for genes examined in this study are as follows: *Tbx6-r.b*, KY21.Chr11.465/466/467; *Hebp-r.a,* KY21.Chr10.215; *Hand-r*, KY21.Chr1.1701; *Tbx6-r.a*, KY21.Chr11.458; *Msx*, KY21.Chr2.1031; *Dlx.b*, KY21.Chr7.361; *Snai*, KY21.Chr3.1356; *Zic-r.b*, KY21.Chr6.26/27/28/29/30/31; *Sox1/2/3*, KY21.Chr1.254; *Lmx1*, KY21.Chr9.606; *Acta.a*, KY21.Chr1.1745; *Tfap2-r.b*, KY21.Chr7.1145; *Pax3/7*, KY21.Chr10.288; *Ets1/2.b*, KY21.Chr10.346; *Id.b*, KY21.Chr7.1129; *Celf3.a,* KY21.Chr6.58; *Slc39a-related*, KY21.Chr4.1089. Identifiers for the latest KY21 set^61^ are shown.

### Functional assays

Sequences of MOs that block translation, were as follows: *Dlx.b,* 5’-TCGGAGATTCAACGACGCTTGACAT-3’; *Zic-r.b*, 5’-GATCAACCATTACATTAGAATACAT-3’; *Snai*, 5’-GTCATGATGTAATCACAGTAATATA-3’; *Msx*, 5’-ATTCGTTTACTGTCATTTTTAATTT-3’. These MOs were used and evaluated in previous studies^8,25,36,39,43,62,63^. They were microinjected under a microscope. For overexpression, coding sequences of *Tbx6-r.b* were cloned into the downstream region of the upstream regulatory region of *Msx* (Chr2:6,132,658-6,133,440). Overexpression constructs were introduced by electroporation^64^. All functional assays were performed at least twice with different batches of embryos. Cells were identified under a microscope according to previous studies (Figure S14)^21,65,66^.

### *In situ* hybridization and reporter assay

Embryos were fixed in 4% paraformaldehyde in 0.1 M MOPS buffer (pH 7.5) and 0.5 M NaCl at 4 °C for over 16 h. After washing with 80% ethanol and phosphate-buffered saline containing 0.1% Tween 20 (PBST), embryos were incubated in 3% H_2_O_2_ for 60 min, and again washed with PBST. Then, embryos were treated with 2 μg/mL proteinase K in PBST for 30 min at 37 °C, and washed with PBST. Embryos were again fixed with 4% paraformaldehyde in PBST for 1 h at room temperature, washed with PBST, and immersed in hybridization buffer for at least 1 h at 50 °C. Then, hybridization buffer was replaced with fresh hybridization buffer containing a digoxigenin- and/or a fluorescein-labeled probe, and embryos were incubated at 50 °C for at least 16 h. Hybridization buffer contained 50% formamide, 5×SSC, 100 μg/mL yeast tRNA, 5×Denhart’s solution, and 0.1% Tween 20. After hybridization, embryos were washed twice at 50 °C for 15 min in 2×SSC, 50% formamide, and 0.1% Tween 20. Embryos were treated for 30 min at 37 °C with 20 μg/mL RNase A in 10 mM Tris buffer (pH 8.0), 0.5 M NaCl, 5 mM EDTA, and 0.1% Tween 20. Embryos were further washed twice at 50 °C for 15 min in 0.5×SSC, 50% formamide, and 0.1% Tween 20, and the washing solution was replaced with PBST. Embryos were blocked at room temperature for 60 min with 0.1% BSA in PBST, and then exposed to mouse anti-digoxigenin antibody (Roche, #11333062910; 1/100 dilution) or rabbit anti-fluorescein antibody (Abcam, ab19491; 1/1000 dilution) diluted in Can-Get-Signal Immunostain Immunoreaction enhancer solution A (Toyobo, NKB-501) at 4 °C overnight. After washing with PBST, embryos were incubated with HRP-conjugated secondary antibody (Thermo Fisher, #B40961 and #B40922). Embryos were washed with PBST and TNT [100 mM Tris buffer (pH7.5), 150 mM NaCl, and 0.1% Tween 20]. Then, embryos were stained with Alexa Fluor 488 or 555 tyramide reagent (Thermo Fisher, #B40953 and #B40955). For two-color *in situ* hybridization, two probes were simultaneously added to the hybridization solution, and the secondary-antibody reaction and detection with tyramide reagents were repeated twice. Before observation with a microscope, embryos were treated with TrueVIEW Autofluorescence Quenching Kit (Vector, #SP-8400-15).

The upstream region that is contained in reporter of *Hebp-r.a>Venus* or *Hebp-r.a>Kaede* is Chr10:1,456,230-1,457,052. The upstream region of *Msx>RFP* that promotes expression in LNPCs and their sibling lineage of cells^67^ is Chr2:6,132,658-6,133,440. These reporter constructs were introduced by electroporation^64^ or microinjection. Expression of fluorescent proteins was examined under fluorescence microscope.

### Immunostaining

For immunostaining of GFP and RFP, embryos were fixed with 3.7% formaldehyde in 0.1 M MOPS buffer (pH 7.5) and 0.5 M NaCl for 5 min at room temperature. After washing with PBSTr (phosphate-buffered saline containing 0.2% Triton-X-100) and PBSTT (phosphate-buffered saline containing 0.2% Triton-X-100 and 0.4% tween 20), embryos were incubated in blocking buffer (10% BSA in PBSTr) for 30 min and then with rabbit and mouse antibodies against GFP (1:250, Thermo Fisher, #A6455) and RFP (1:250, Medical & biological laboratories, #M1553) for 6 h at 4 °C. Then, embryos were washed with PBSTr, and incubated with secondary antibodies (Alexa Fluor 488-conjugated anti-rabbit antibody, 1:400, Thermo Fisher, #A21206; Alexa Fluor 555-conjugated anti-mouse antibody, 1:400, Thermo Fisher, #A31570) for 6 h at 4 °C. Embryos were washed with PBSTr and mounted on slide glass using Vectashield mounting medium with DAPI (vector laboratories, H-1200).

### Data analysis

To compare gene expression profiles among *Ciona* larval tissues, we used single-cell transcriptome data for *Ciona* larvae that were published previously^34^ (SRA accession numbers: SRR9051005 to SRR9051007). Data were mapped to the latest genome assembly (HT version)^68^ and the latest version of the gene model set (KY21 version)^61^ using Cell Ranger software (version 6.12. 10xGenomics). Then, mesenchymal cells, notochord cells, muscle cells, endodermal cells, endodermal strand cells, tail-tip cells, nerve cord cells, brain cells, sensory neurons, other neural cells, and epidermal cells were annotated using Loupe Browser (10xGenomics) (Table S6). For cell identification during the larval stage, we used *Hebp-r.a*, *Hand-r*, KY21.Chr8.1029 for tail-tip cells, KY21.Chr1.1899 and KY21.Chr1.1498 for mesenchyme^69,70^, KY21.Chr6.610 (*Noto1*)^71^ for notochord, KY21.Chr 11.773 (*Myh.f*) and *Acta.a* for muscle^72^, KY21.Chr1.2012 (*Epi1*) and KY21.Chr7.872 (*Epib*) for epidermal cells^73^, KY21.Chr9.123 (*Gnrh2*) and KY21.Chr10.869 for nerve cord^70,74^, KY21.Chr11.800 (*Rlbp1*) for brain^75^, and KY21.Chr2.456 (*Pou4*) for sensory neurons^30^. Other neural cells were identified by expression of KY21.Chr11.1137 and KY21.Chr2.49^34^, KY21.Chr6.690 (*Pax2/5/8.a*)^36^, or KY21.Chr2.1203 (*Syt*)^76^. Endodermal cells were identified by expression of KY21.Chr6.222 (*Alp*) and KY21.Chr14.805^32^, and endodermal strand cells were identified as cells expressing KY21.Chr14.805, without expression of *Alp*^32,77^. A heatmap containing the top 10 genes for each of 11 tissues was generated with Seurat^78^ (version 4.1.3) as follows. Expression matrices of three larval samples were loaded into three Seurat objects. We normalized each dataset using the ‘NormalizeData’ function with default parameters, and calculated 800 highly variable features using the Seurat ‘FindVariableFeatures’ function with default parameters on the basis of plots obtained with the ‘VariableFeaturesPlot’ function. A list of three Seurat objects was used for selecting features that are variable across these datasets using the ‘SelectIntegrationFeatures’ function with default parameters. Then, the ‘FindIntegrationAnchors’ function was used to identify anchors for data integration with default parameters. Using these anchors, the above eight datasets were integrated using the ‘IntegrateData’ function with default parameters except ‘k.weight = 10’. The assay mode was set to ‘integrated’ with the ‘DefaultAssay’ function, and data were scaled with the ‘ScaleData’ function. We chose only cells that we annotated as mesenchymal cells, notochord cells, muscle cells, endodermal cells, endodermal strand cells, tail-tip cells, nerve cord cells, brain cells, sensory neurons, other neural cells, or epidermal cells, as mentioned above. Then, we tried to find genes differentially expressed in each of these cell types using the ‘FindAllMarkers’ function with default parameters except ‘only.pos=TRUE’ and ‘min.pct=0.25’. Then we picked top the 10 genes for each of these cell types. After calculating averages among cells with the ‘AverageExpression’ function, we created a heatmap with ‘DoHeatmap’ function.

For a cross-species comparison between *Ciona* transcriptomes and zebrafish transcriptomes, single-cell transcriptome data for *Ciona* middle gastrula embryos to larvae, which were published previously^34^, were used (SRA accession numbers: SRR9050988, SRR9050989, SRR9050992, SRR9050997, SRR9050998, SRR9051003, and SRR9051007). Data were mapped to the latest genome assembly (HT version)^68^ and the latest version of the gene model set (KY21 version)^61^ using Cell Ranger software (version 6.12. 10xGenomics). For zebrafish 10- to 24-hr embryos, mapping data to gene models were downloaded from the GEO database (we used CSV files of UMI-filtered counts under accession numbers, GSM3067192 to GSM3067195; GSM3067192_10hpf.csv.gz, GSM3067193_14hpf.csv.gz, GSM3067194_18hpf.csv.gz, and GSM3067195_24hpf.csv.gz)^44^.

Annotations of zebrafish cells published previously^44^ were used. We annotated *Ciona* cells using Loupe Browser (10xGenomics). Tail-tip cells, muscle and notochord cells in middle tailbud embryos were identified using the same marker genes that were used in larvae. At the middle gastrula and early neurula stages, we identified b9.34 as cells expressing *Msx*, *Tbx6-r.b*, and *Snai*, and b9.33 as cells expressing *Msx*, *Snai*, and *Hox12* but not *Pax3/7* or *Tbx6-r.b*. These cells were further curated manually and are listed in Table S7.

For comparisons with zebrafish transcriptomes, we first compared proteomes of *Ciona* (KY21 version) and zebrafish (Refseq^79^ proteome file named GCF_000002035.6_GRCz11_protein.faa) using OrthoFinder^45^. We took 10 protein groups from the *Ciona* proteome to validate grouping by OrthoFinder, and found that comparisons between *Ciona* and zebrafish proteomes, which yielded 5422 groups, were not satisfactory (Table S8). According to the manual of Orthofinder, we included proteomes of *Strongylocentrotus purpuratus* (sea urchin) and *Drosophila melanogaster* as controls to obtain better resolution. Proteome files we used were: “HT.KY21Gene.protein.2.fasta.zip” for *Ciona*, which includes a proteome set derived from the latest KY21 gene model set^61^, downloaded from the Ghost database^31^; “GRCz11_protein.faa” for the zebrafish proteome, downloaded from the NCBI Refseq database^79^; “GCF_000002235.5_Spur_5.0_translated_cds.faa.gz” was downloaded for the sea urchin proteome from the NCBI Refseq database^79^; and “Drosophila_melanogaster.BDGP6.32.pep.all.fa.gz” for the fly proteome, downloaded from the Ensembl database (release 108). From these data, OrthoFinder yielded 6379 ortholog groups that contained both ascidian and zebrafish proteins (Table S9 and Table S10). We confirmed that proteins in the above control 10 groups are resolved better (Table S8). When multiple genes for one species were included in one ortholog group, all read counts in that group were summed.

Data were analyzed with Seurat^78^ (version 4.1.3). After converting all gene names to ortholog-group IDs, zebrafish data were loaded into a single Seurat^78^ object (listed in Table S3; 13,951 cells). We normalized the zebrafish data using the ‘NormalizeData’ function with default parameters, and calculated 800 highly variable features using the Seurat ‘FindVariableFeatures’ function with default parameters on the basis of a plot obtained with the ‘VariableFeaturesPlot’ function. *Ciona* expression matrices were individually loaded into seven Seurat objects, because these were not normalized against one another (middle gastrula, 2296 cells; early neurula, 1916 cells; late neurula, 6177 cells; middle tailbud II replica 1, 5233 cells; middle tailbud II replica 2, 4329 cells; late tailbud II, 5277 cells; larva, 6459 cells). On the basis of scatter plots created with the ‘FeatureScatter’ function, we chose cells that have unique counts within the ranges shown in Table S11 using the ‘subset’ function (middle gastrula, 1727 cells; early neurula, 1309 cells; late neurula, 1600 cells; middle tailbud II replica 1, 2254 cells; middle tailbud II replica 2, 1853 cells; late tailbud II, 2221 cells; larva, 4007 cells). We normalized the data using the ‘NomalizeData’ function with default parameters, and calculated 300 highly variable features similarly (this value was determined on the basis of plots obtained with the ‘VariableFeaturesPlot’ function). A list of seven *Ciona* Seurat objects and one zebrafish Seurat object was used for selecting features that are variable across these datasets using the ‘SelectIntegrationFeatures’ function with default parameters. Then, the ‘FindIntegrationAnchors’ function was used to identify anchors for data integration with default parameters. Using these anchors, the above eight datasets were integrated using the ‘IntegrateData’ function with default parameters except ‘k.weight = 10’. From the integrated object, we removed irrelevant *Ciona* cells (cells other than b9.33, b9.34, b9.37, b9.38, muscle, notochord, and tail-tip cells; the remaining 301 cells are listed in Table S7). Note that we did not remove any zebrafish cells at this stage. The assay mode was set to ‘integrated’ with the ‘DefaultAssay’ function. After scaling the data with the ‘ScaleData’ function, a principal component analysis was performed with the ‘RunPCA’ function with default parameters. To cluster cells, we used the ‘FindNeighbors’ function with the parameter ‘dims=1:50’, and the ‘FindCluster’ function with the parameter ‘resolution=10’ (high resolution), ‘resolution=0.25’ (middle resolution), and ‘resolution=0.2’ (low resolution). Data were visualized with the ‘RunUMAP’ function with default parameters.

All matrices used for the transcriptome comparison between *Ciona* and zebrafish are shown as Supplementary Data S1-S8.

---

**Figure S1.**
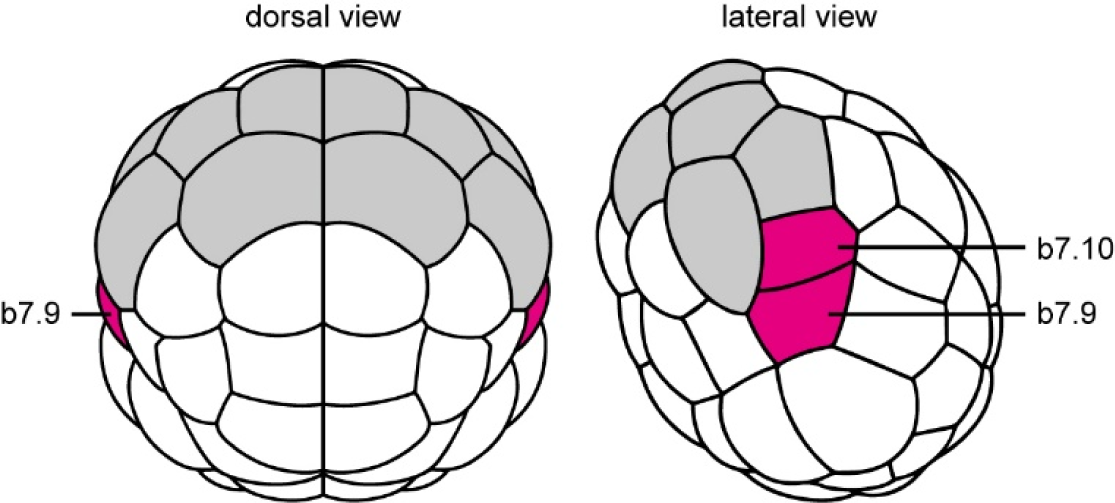
An ascidian 64-cell embryo. Cells that contribute to the central nervous system are colored. b7.9 and b7.10, which produce LNPCs, are colored magenta, and other cells that give rise to the neural plate are colored gray.

**Figure S2.**
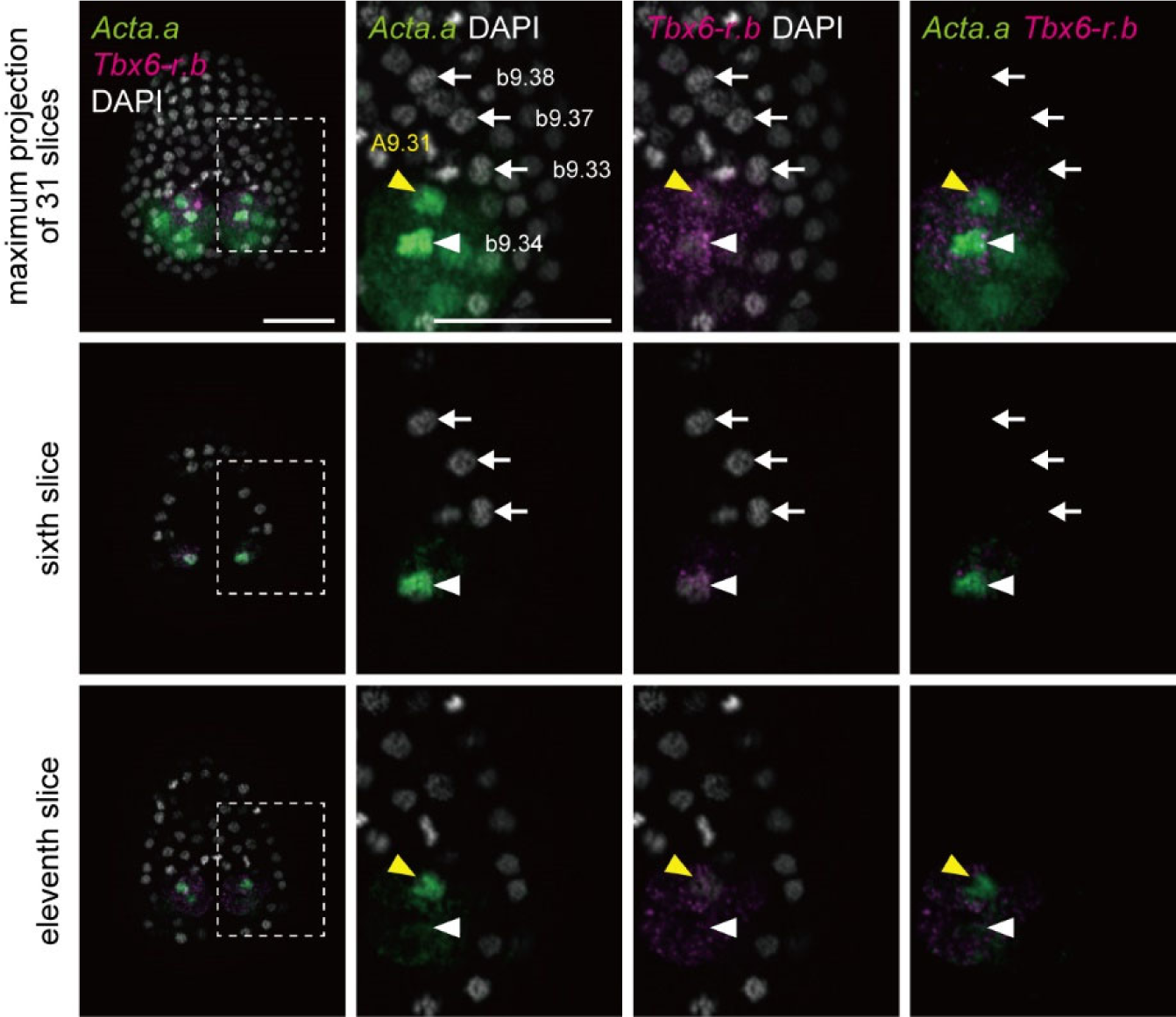
*Acta.a*, which encodes a muscle actin, is expressed in b9.34 at the late gastrula stage. *Acta.a* expression was examined by *in situ* hybridization (green). *Tbx6-r.b* expression was also examined (magenta). Z-projected image stacks overlaid in pseudocolor are shown in the top row. The sixth optical slice more clearly shows that *Acta.a* and *Tbx6-r.b* are expressed in b9.34, but not in the other LNPCs. The eleventh optical slice shows expression of *Acta.a* and *Tbx6-r.b* in A9.31. Brightness and contrast of photographs were linearly adjusted.

**Figure S3.**
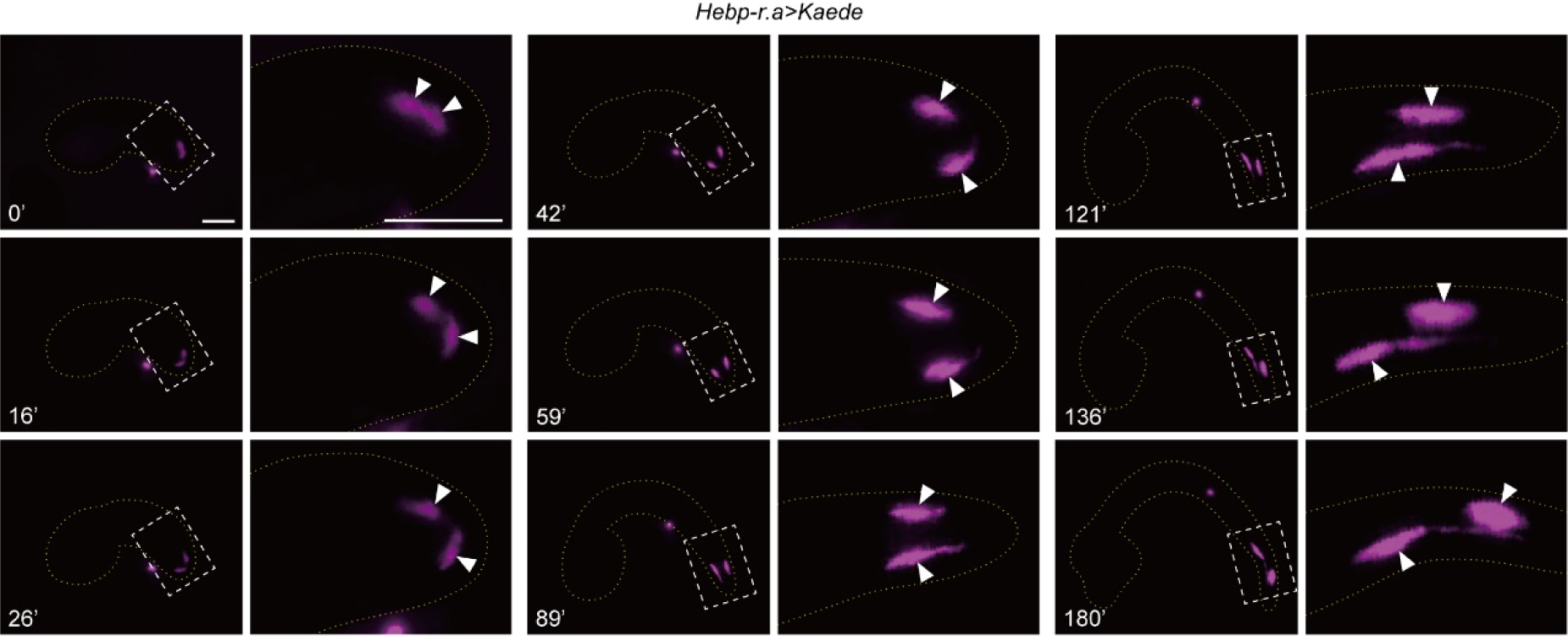
Some pLNPC-derived cells change location from the dorsal to the ventral side between early and late tailbud stages. A tailbud embryo expressing *Hebp-r.a>Kaede* reporter was UV-irradiated for photoconversion of Kaede fluorescence at the early tailbud stage (0 min; early tailbud I). One cell with photoconverted Kaede changed its location to the ventral side. Note that only two tail-tip cells are labelled because of mosaic incorporation of the reporter construct. The first, third, and sixth photographs are the same as the photograph shown in Figure 1e. Photographs are z-projected image stacks overlaid in pseudocolor. Brightness and contrast of photographs were linearly adjusted. The dorsal side is up, and the ventral side is down.

**Figure S4.**
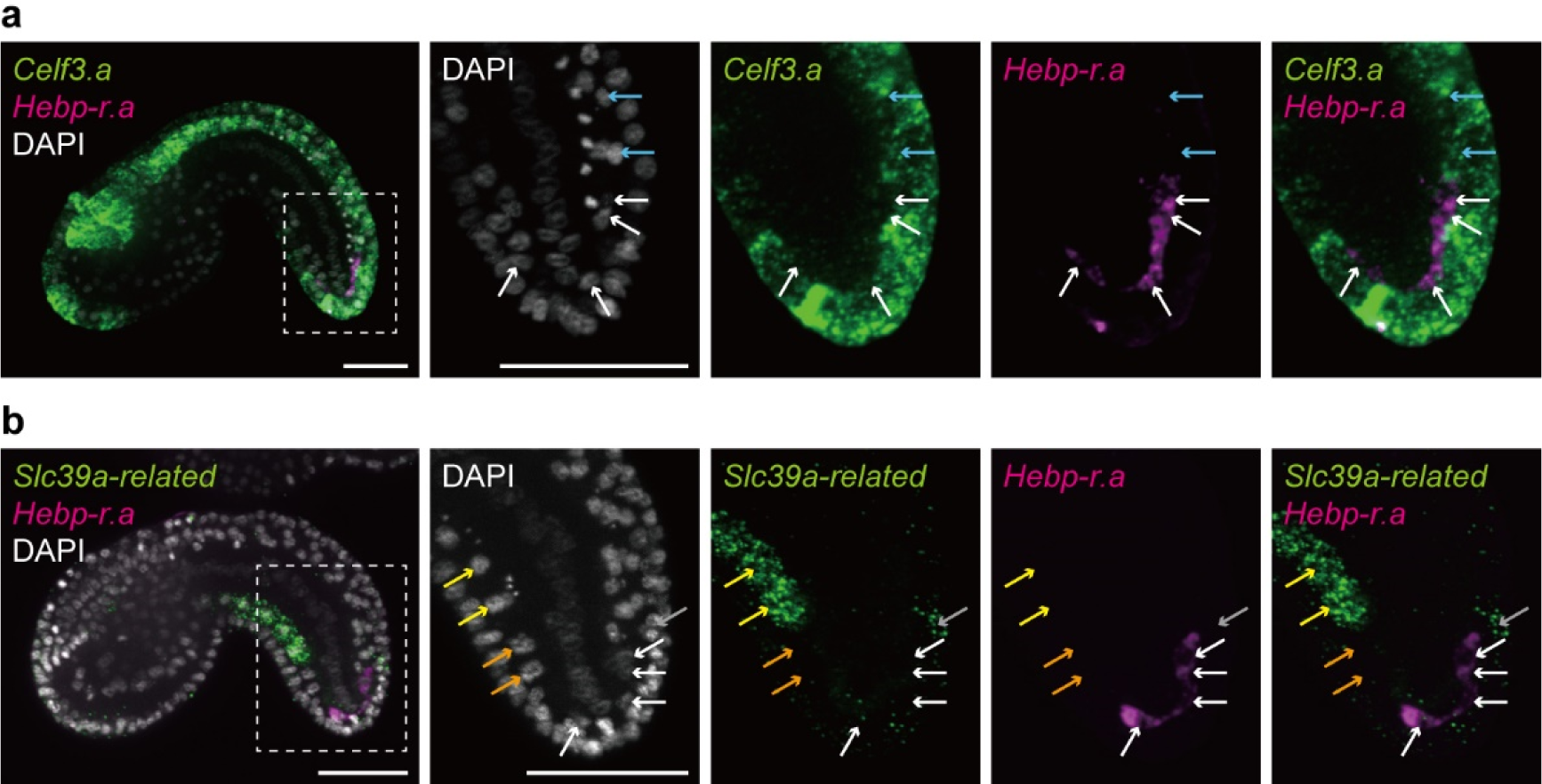
Expression of a pan-neural marker, *Celf3.a*, and an endodermal-strand marker, *Slc39a-related*, in tailbud embryos. Expression was examined by *in situ* hybridization, and photographs are z-projected image stacks overlaid in pseudocolor (*Celf3.a* and *Slc39a-related*, green; *Hebp-r.a*, magenta). Tail-tip cells, which express *Hebp-r.a*, do not express *Celf3.a* or *Slc39a-related*. Brightness and contrast of photographs were linearly adjusted. Tail-tip cells are indicated by white arrows. Nerve cord cells are indicated by cyan arrows, and endodermal strand cells are indicated by yellow arrows. There are two putative germ-line cells between anterior endodermal strand cells and tail-tip cells (orange arrows). Gray arrows indicate epidermal cells. Nuclei are stained with DAPI (gray). Scale bar, 50 μm.

**Figure S5.**
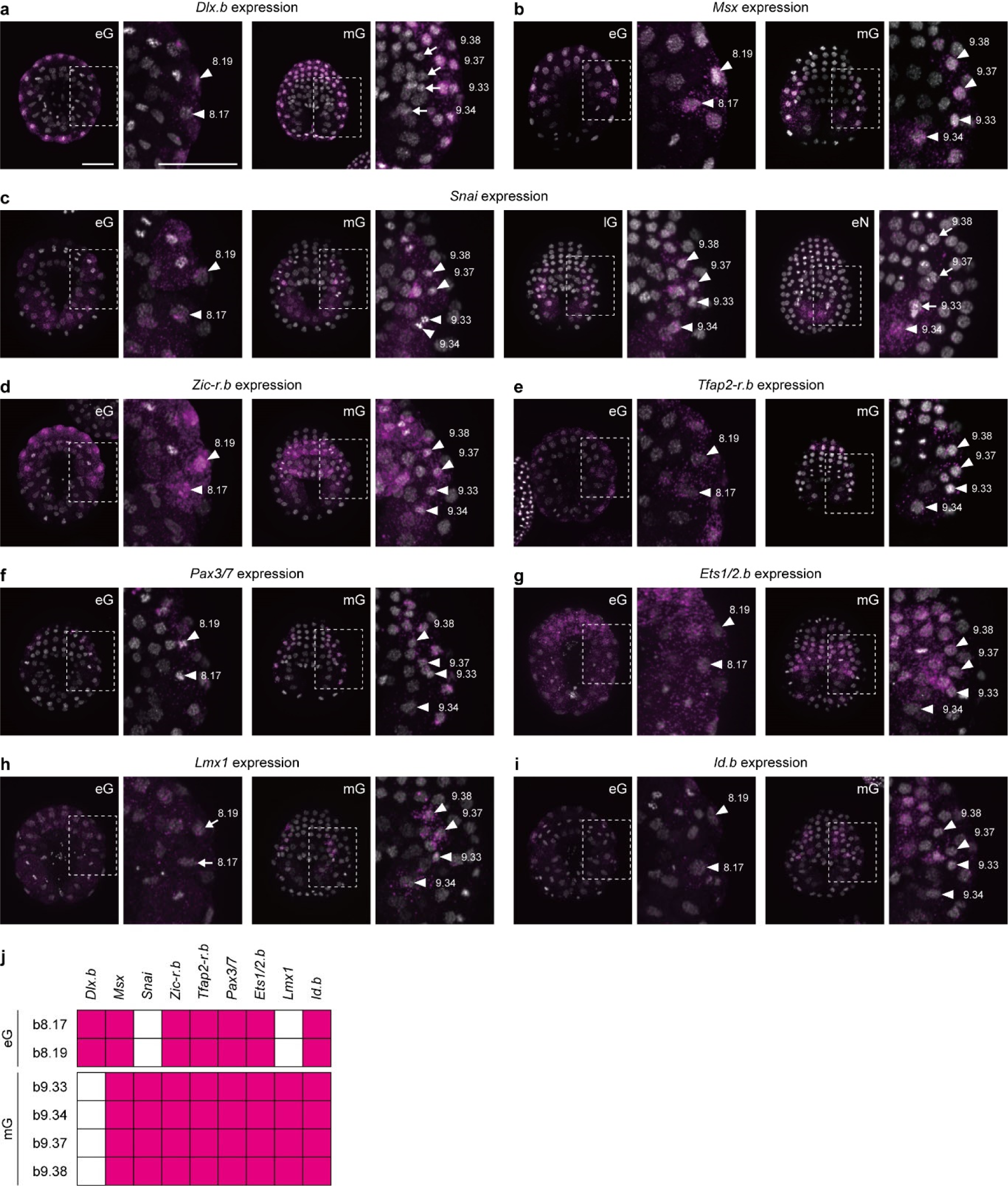
*Dlx.b, Msx*, *Snai*, *Zic-r.b*, *Tfap2-r.b*, *Pax3/7*, *Ets1/2.b*, *Lmx1*, and *Id.b* are expressed in LNPCs. (a-i) Expression was examined by *in situ* hybridization, and photographs are z-projected image stacks and overlaid in pseudocolor (magenta). Nuclei were stained with DAPI (gray). Higher magnification views are shown on the right. Arrowheads indicate gene expression and arrows indicate the absence of expression. Developmental stages are shown in photographs; eG, early gastrula; mG, middle gastrula; eN, early neurula. Scale bar, 50 μm. (j) A summary of gene expression in LNPCs at the early and middle gastrula stages.

**Figure S6.**
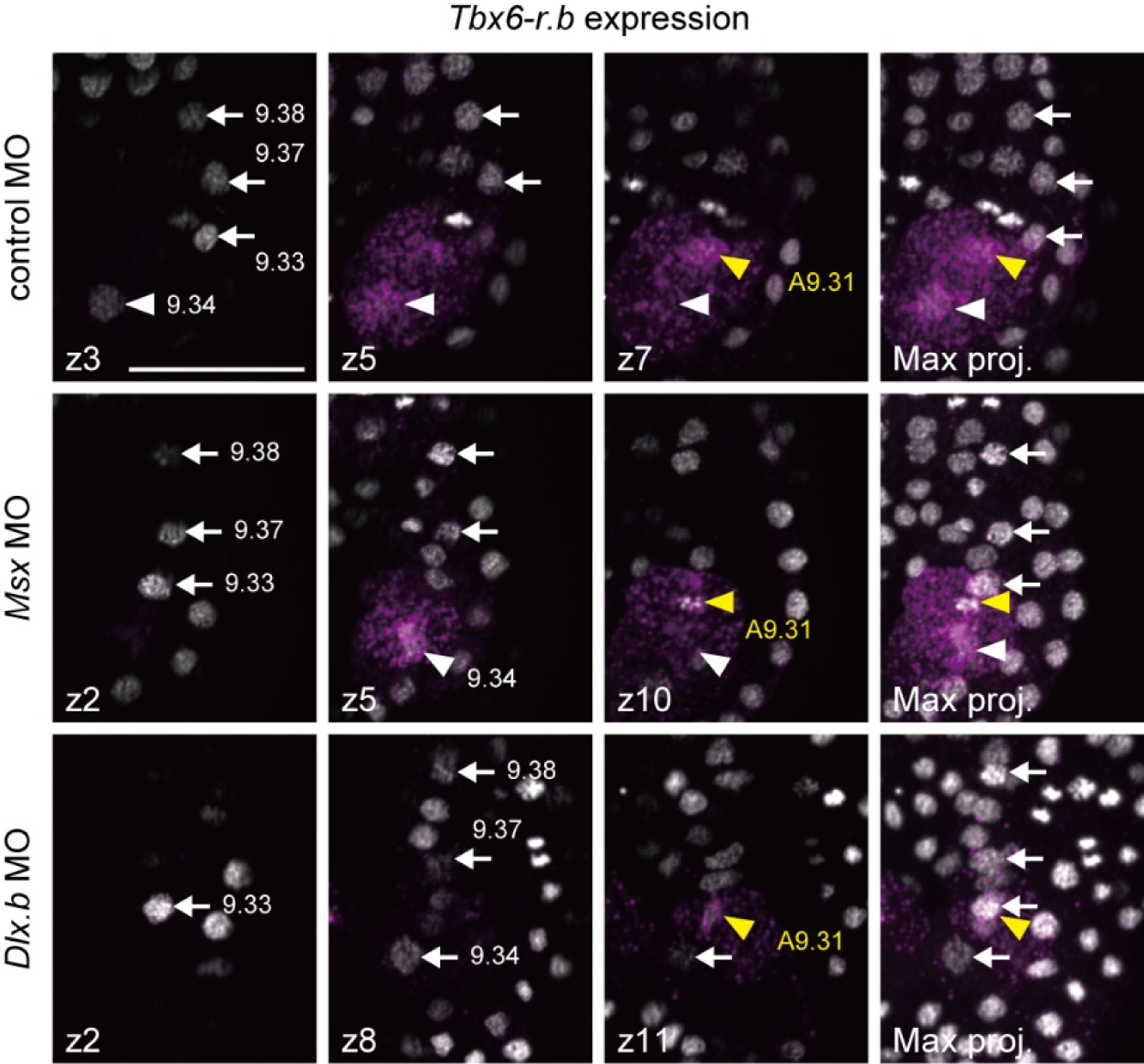
Expression of *Tbx6-r.b* in a control embryo and embryos injected with an MO against *Msx* or *Dlx.b*. Optical slices showing *Tbx6-r.b* expression in control and embryos injected with the *Msx* MO or *Dlx.b* MO are shown in the left three columns, and z-projected image stacks are shown on the right. Z-projected image stacks are the same as the photograph shown in Figure 2. Arrowheads indicate gene expression and arrows indicate the absence of *Tbx6-r.b* expression. Although signals are seen around b9.33 in z-projected image stacks, this signal shows expression in an A-line muscle cell (A9.31; yellow arrowheads), but not in b9.33. Nuclei are stained with DAPI (gray). Brightness and contrast of photographs were linearly adjusted. Photographs are overlaid in pseudocolor. Scale bar, 25 μm.

**Figure S7.**
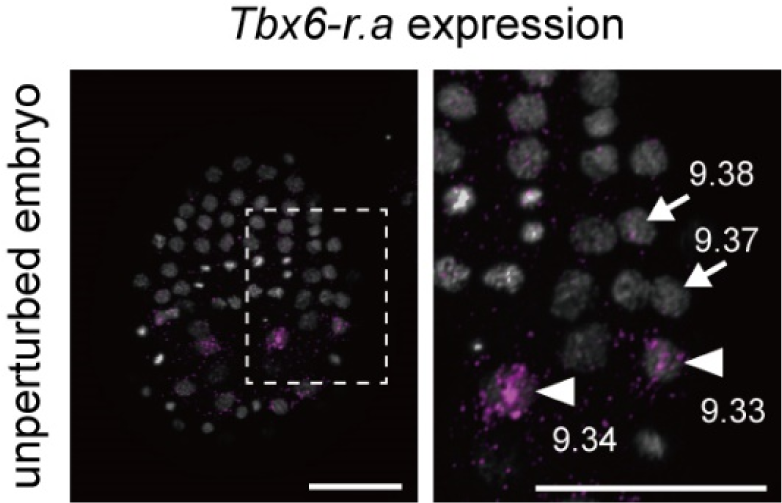
*Tbx6-r.a* expression in a middle gastrula embryo. *Tbx6-r.a* is expressed in pLNPCs (b9.33 and b9.34), and indicated by white arrowheads. The remaining LNPCs (aLNPCs) are shown by arrows. Photographs are z-projected image stacks overlaid in pseudocolor. Nuclei are stained with DAPI (gray), and a higher magnification view is shown on the right. Brightness and contrast of photographs were linearly adjusted. Scale bars, 50 μm.

**Figure S8.**
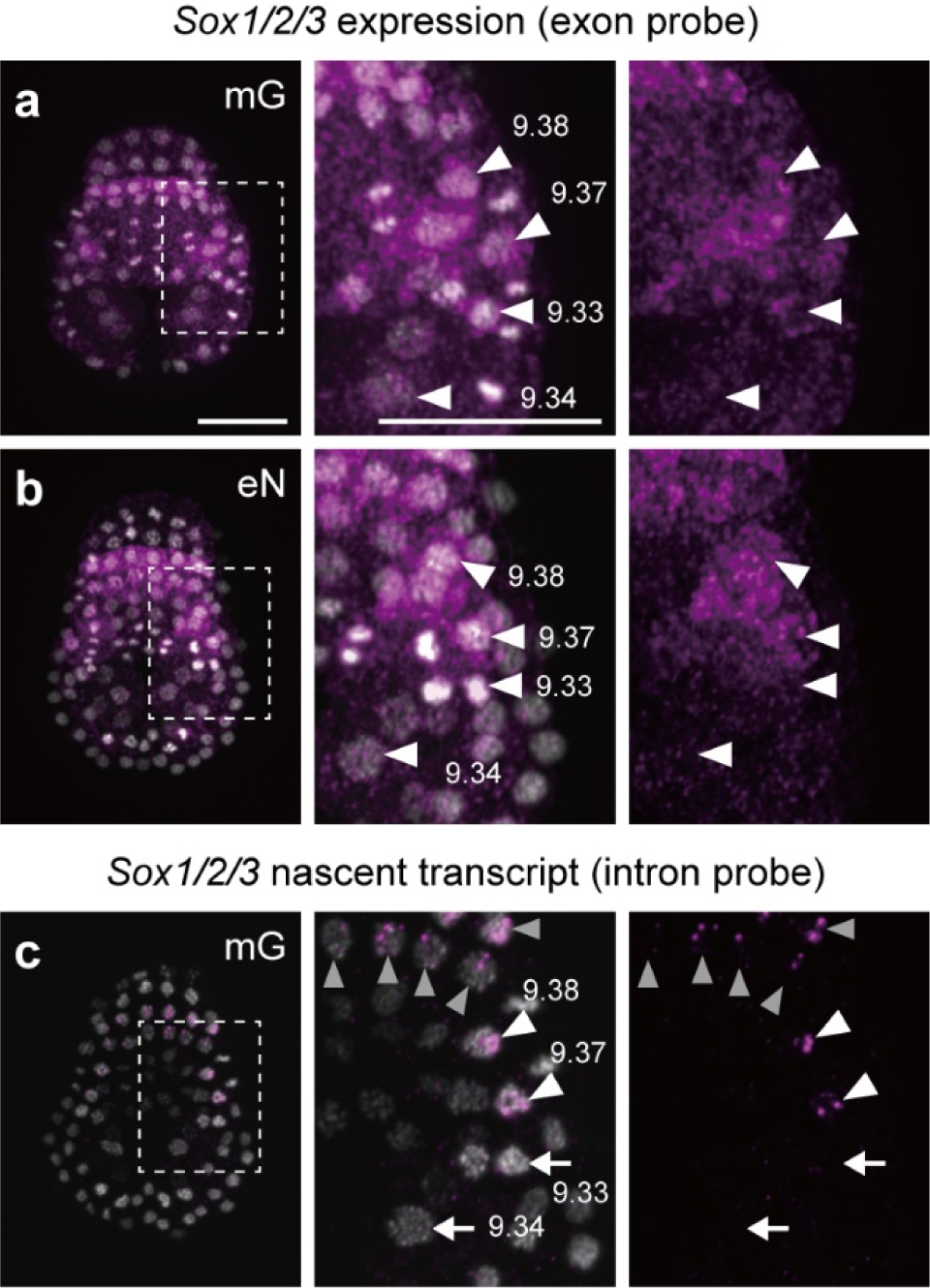
Expression of *Sox1/2/3* in unperturbed middle gastrula embryos. (a, b) *Sox1/2/3* mRNA is detected in cells including LNPCs (white arrowheads) of normal middle gastrula gastrula (mG) and early neurula (eN) embryos. Note that signals in pLNPCs (b9.33 and b9.34) are weak at the early neurula stage. (c) *Sox1/2/3* nascent transcripts, which were examined with an intron probe, are not seen in pLNPCs (white arrows), which indicates that *Sox1/2/3* is not transcribed in pLNPCs. Note that *Sox1/2/3* is expressed in cells that contribute to the central nervous system (gray arrowheads) and aLNPCs (white arrowheads). Photographs are z-projected image stacks overlaid in pseudocolor; magenta, *in situ* hybridization signals; gray, nuclei stained with DAPI. Brightness and contrast of photographs were linearly adjusted. Scale bar, 50 μm.

**Figure S9.**
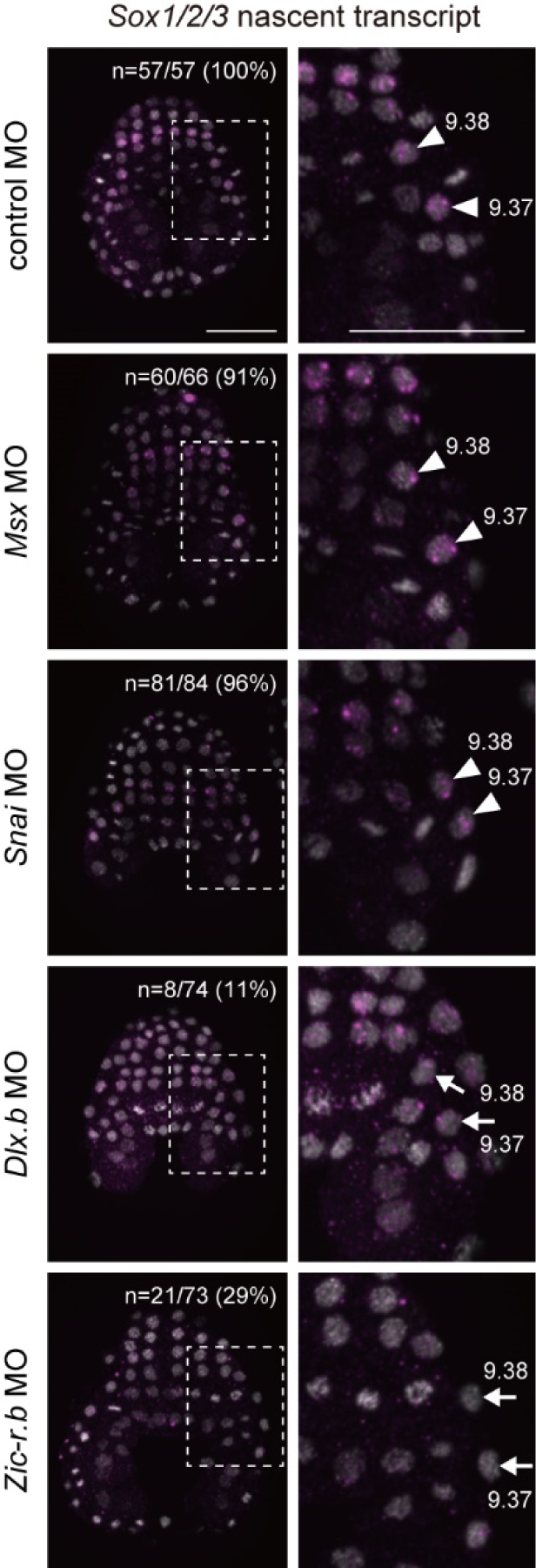
Expression of *Sox1/2/3* in morphant embryos of *Msx*, *Snai*, *Dlx.b*, or *Zic-r.b* at the middle gastrula stage. Expression of *Sox1/2/3* nascent transcripts was examined by *in situ* hybridization using a probe designed to hybridize with the first intron of *Sox1/2/3*. Higher magnification views are shown on the right. LNPCs that express or do not express designated genes are shown by arrowheads and arrows, respectively. Nuclei are stained with DAPI and are shown in gray. Photographs are z-projected image stacks overlaid in pseudocolor. Brightness and contrast of photographs were linearly adjusted. Numbers of embryos examined and embryos that expressed *Sox1/2/3* nascent transcripts in b9.37 and b9.38 are shown in the panels. Scale bars, 50 μm.

**Figure S10.**
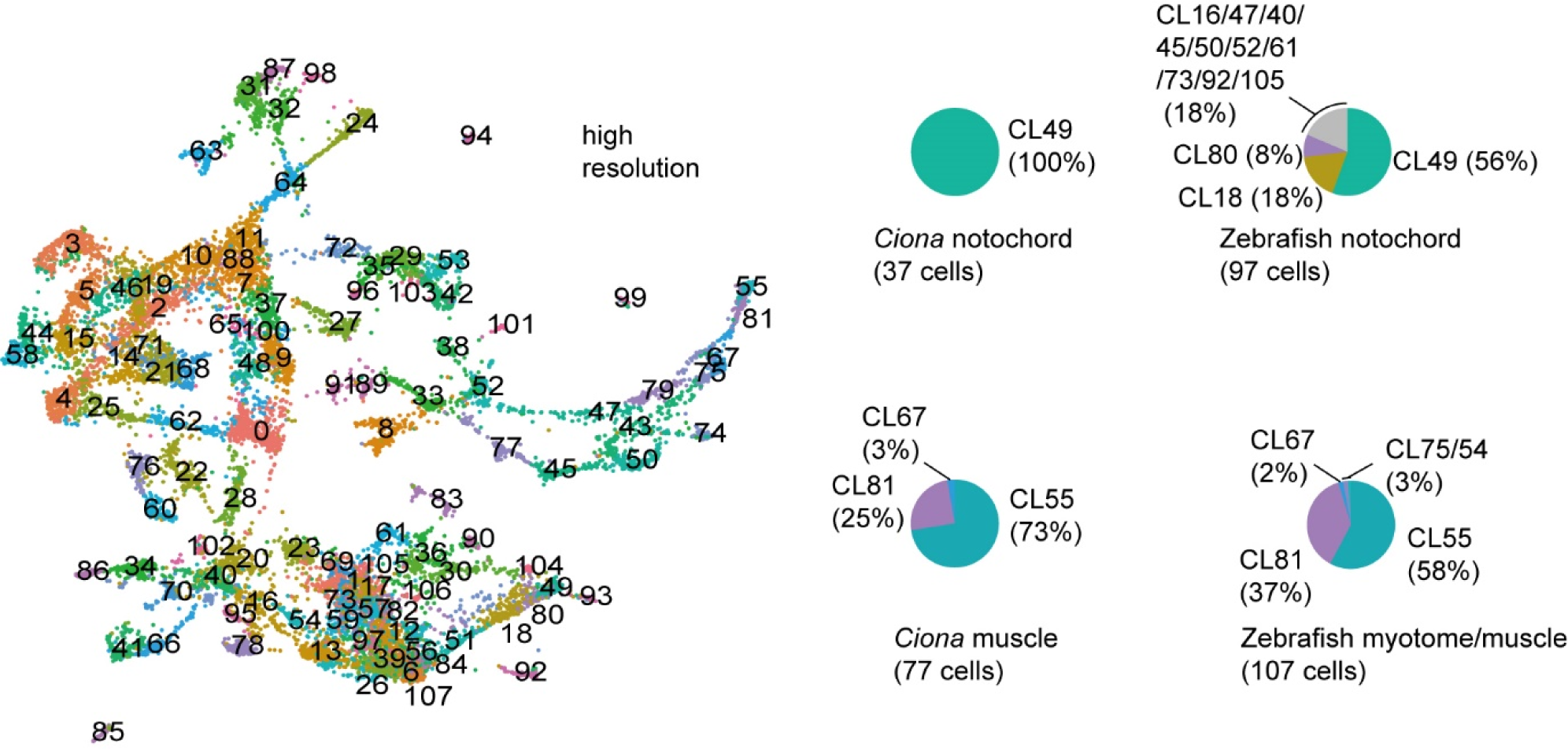
High-resolution clustering of single-cell transcriptome data of *Ciona* and zebrafish. High-resolution clustering results are mapped on the same UMAP plot that is shown in Figure 4. Different clusters are indicated by different colors. Pie charts show that notochord and muscle cells of *Ciona* and zebrafish are largely grouped into the same clusters.

**Figure S11.**
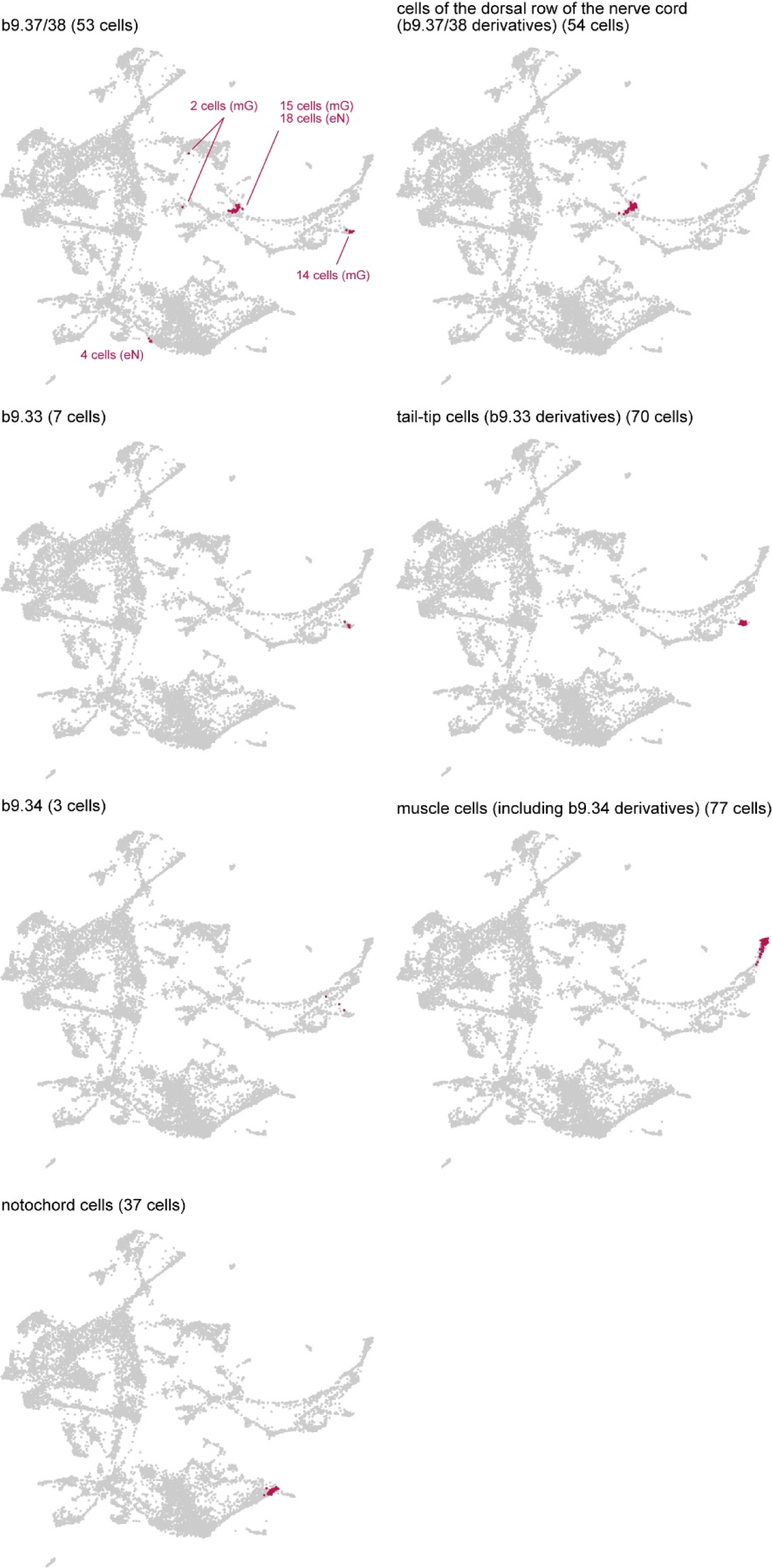
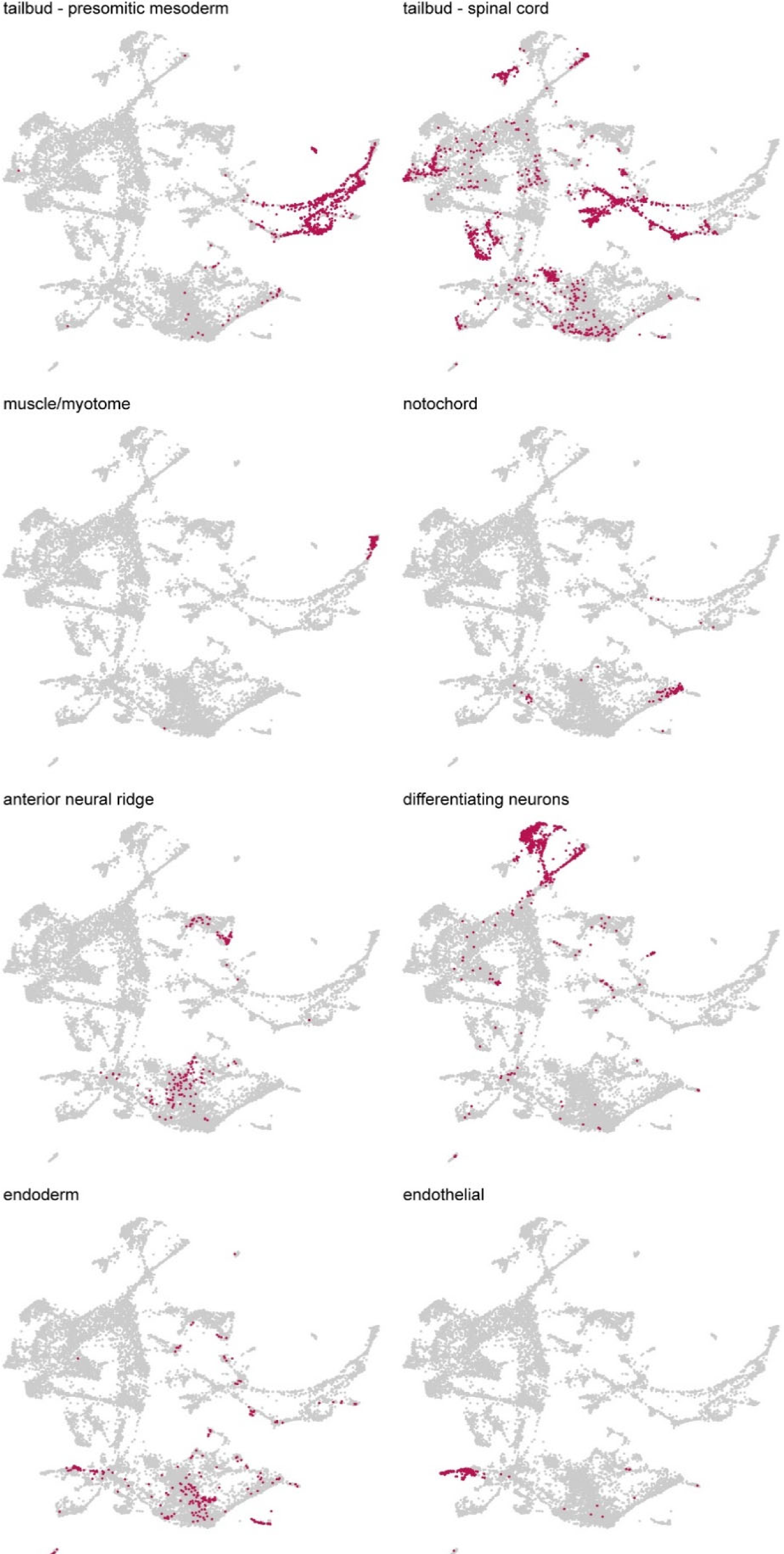
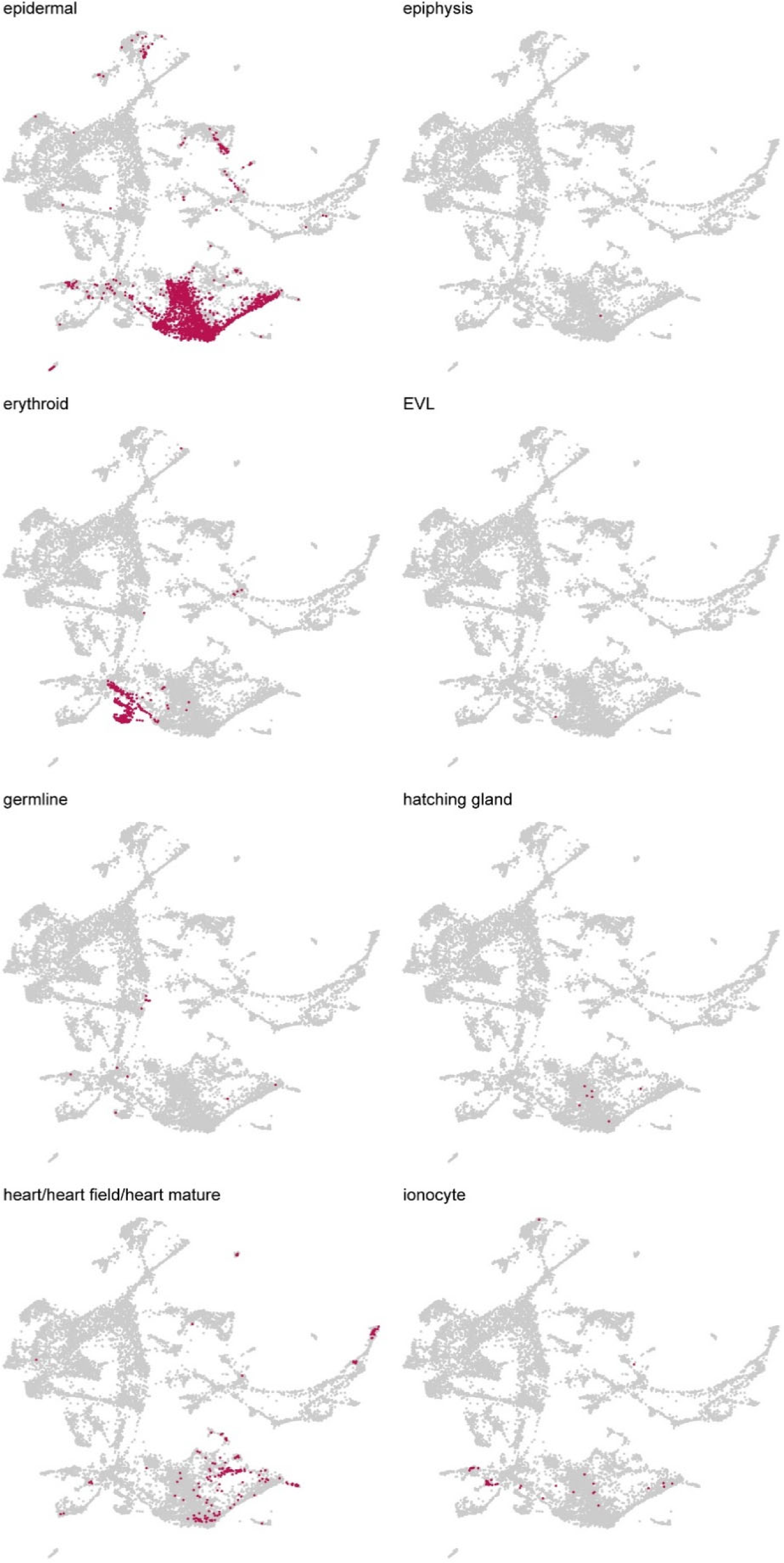
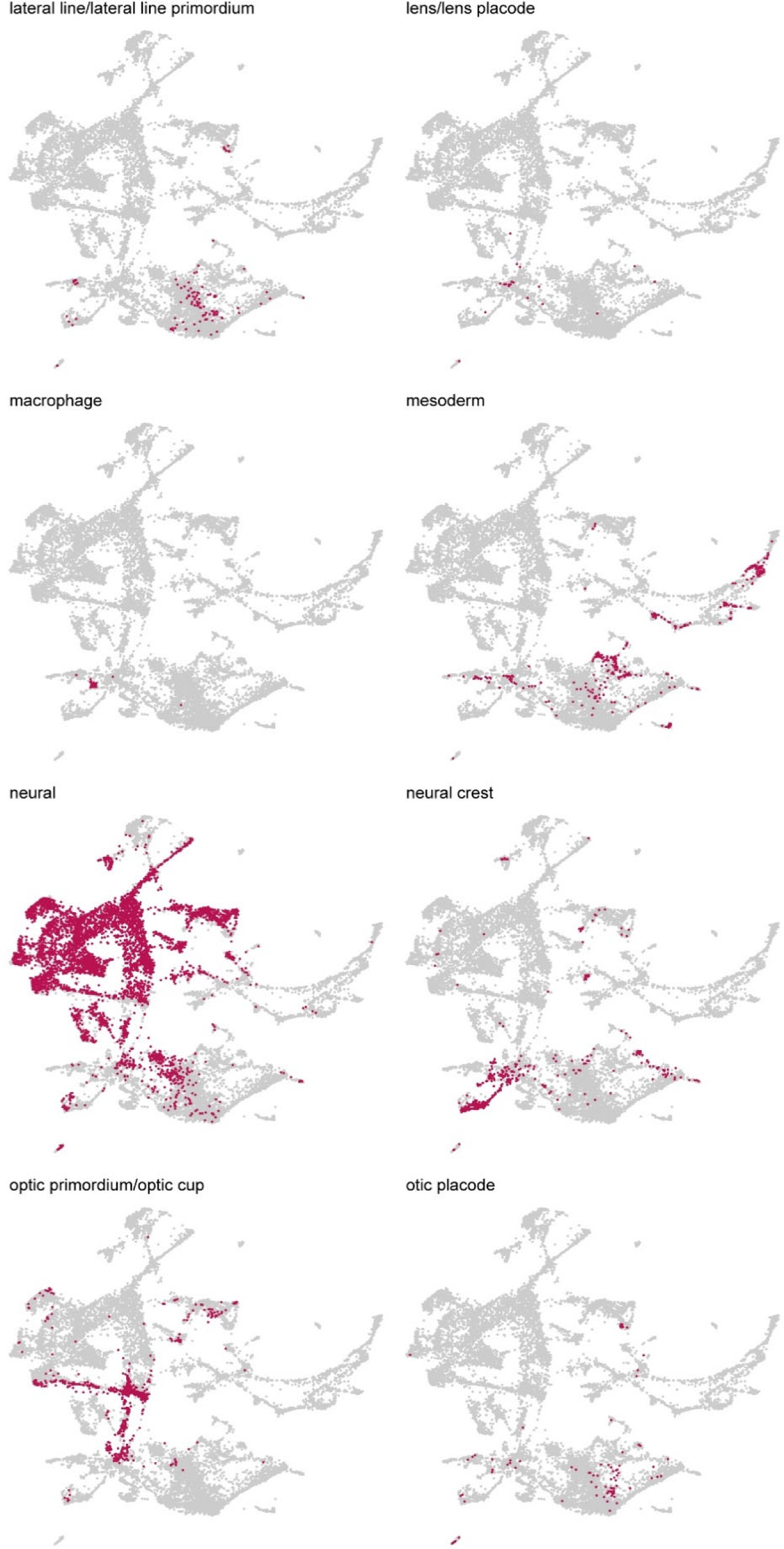
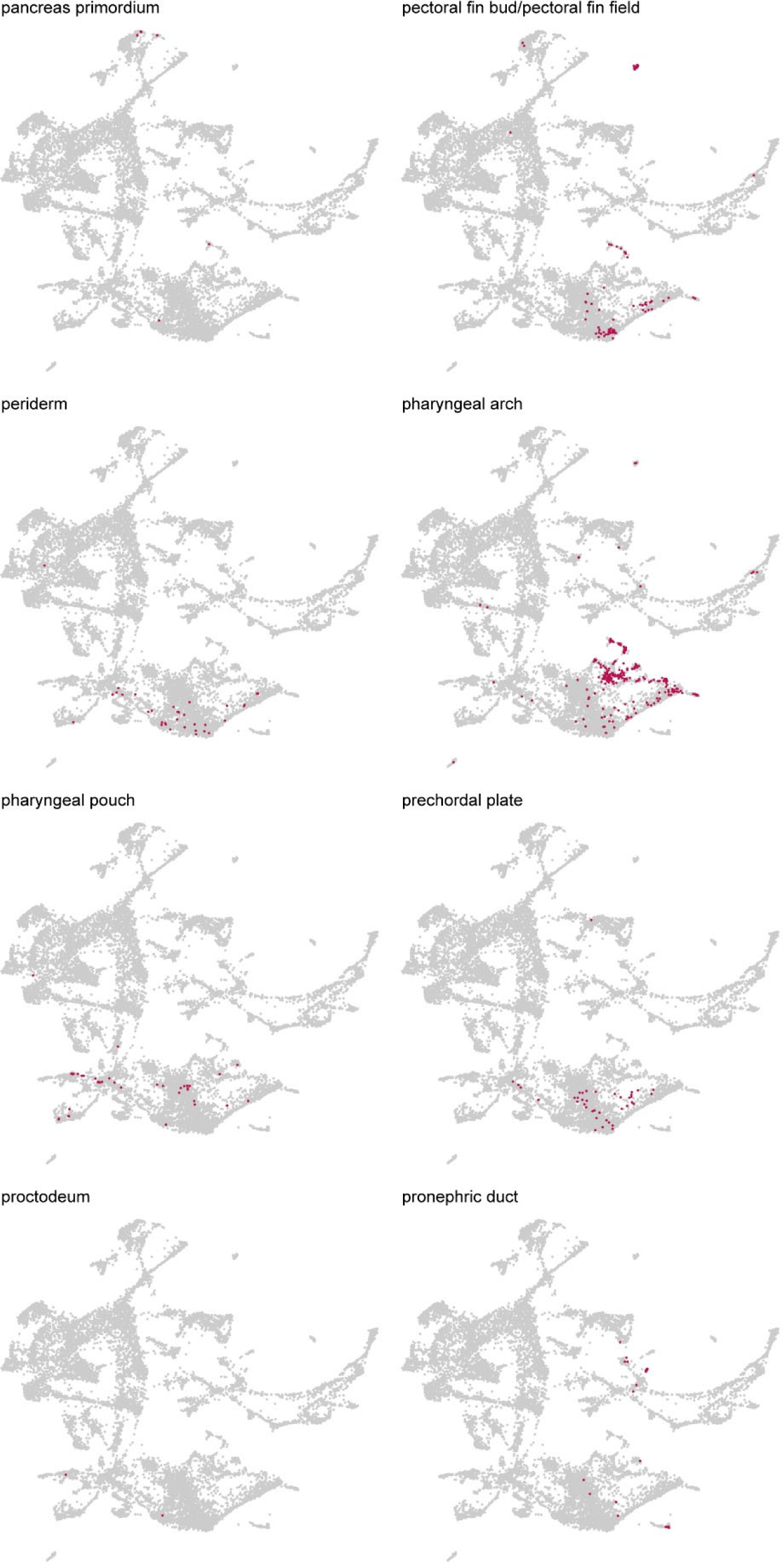
Locations of individual cell types of *Ciona* in the cross-species comparison UMAP shown in Figure 4. Seven types of cell in *Ciona* embryos are individually shown by magenta dots indicating cells.

**Figure S12.**
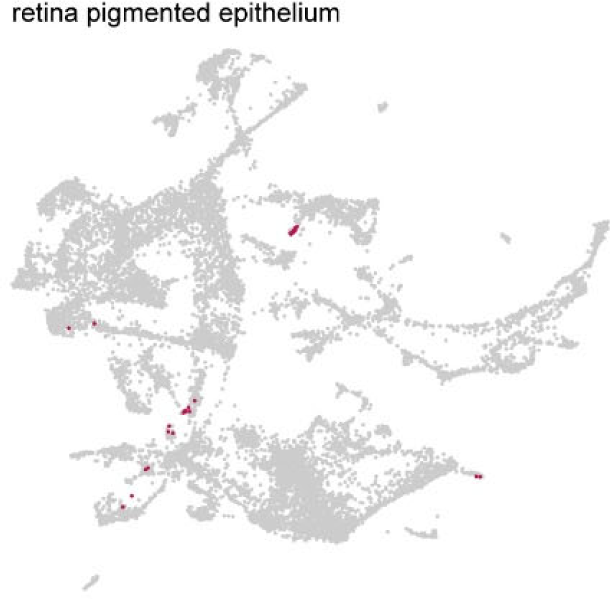
Locations of individual cell types of zebrafish in the cross-species comparison UMAP shown in Figure 4. Cell types in zebrafish embryos are individually shown by magenta dots indicating cells.

**Figure S13.**
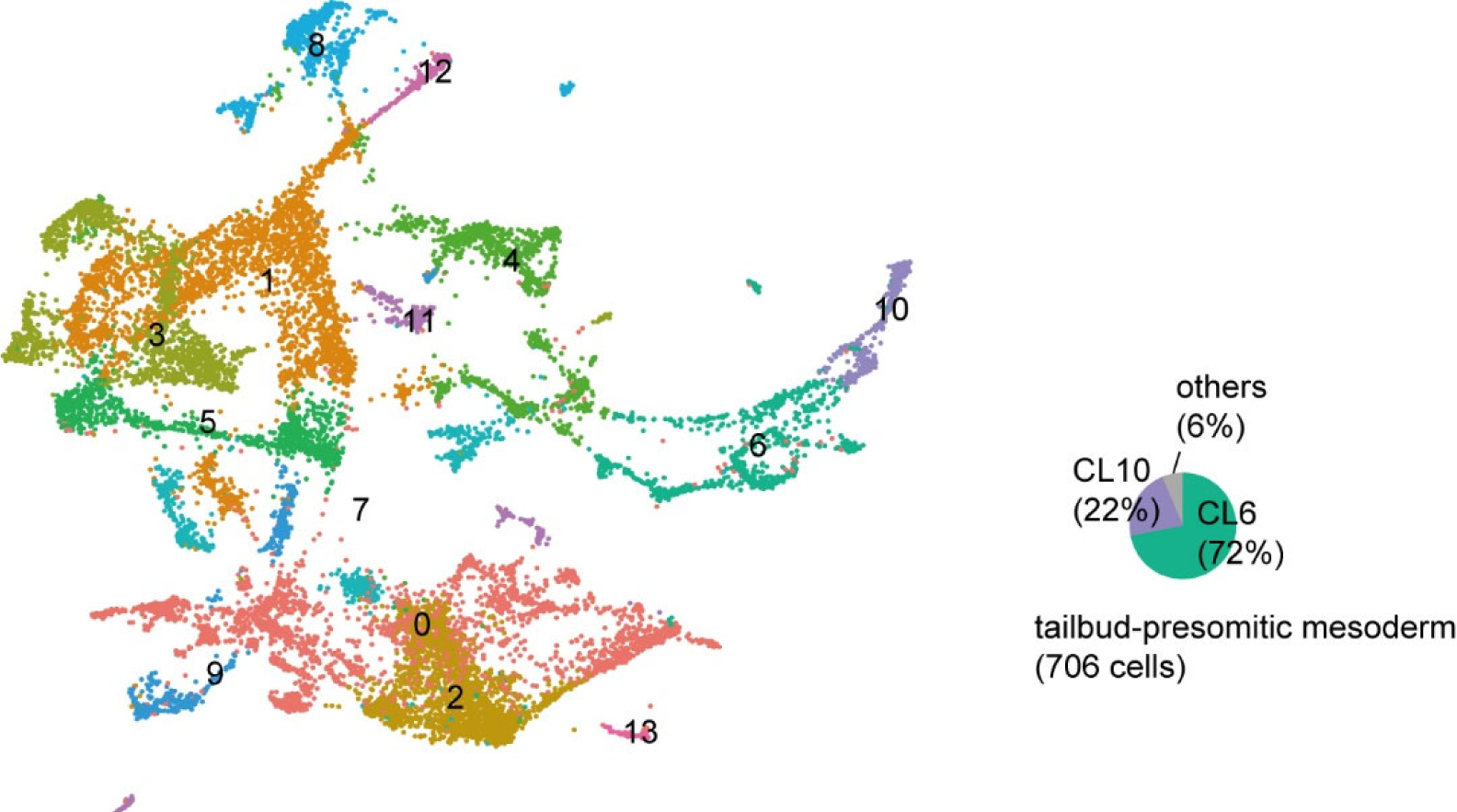
Low-resolution clustering of single-cell transcriptome data of *Ciona* and zebrafish. Low-resolution clustering results are mapped on the same UMAP plot that is shown in Figure 4. Different clusters are indicated by different colors. The pie chart shows that zebrafish cells annotated “tailbud-presomitic mesoderm” are mostly in clusters 6 and 10.

**Figure S14.**
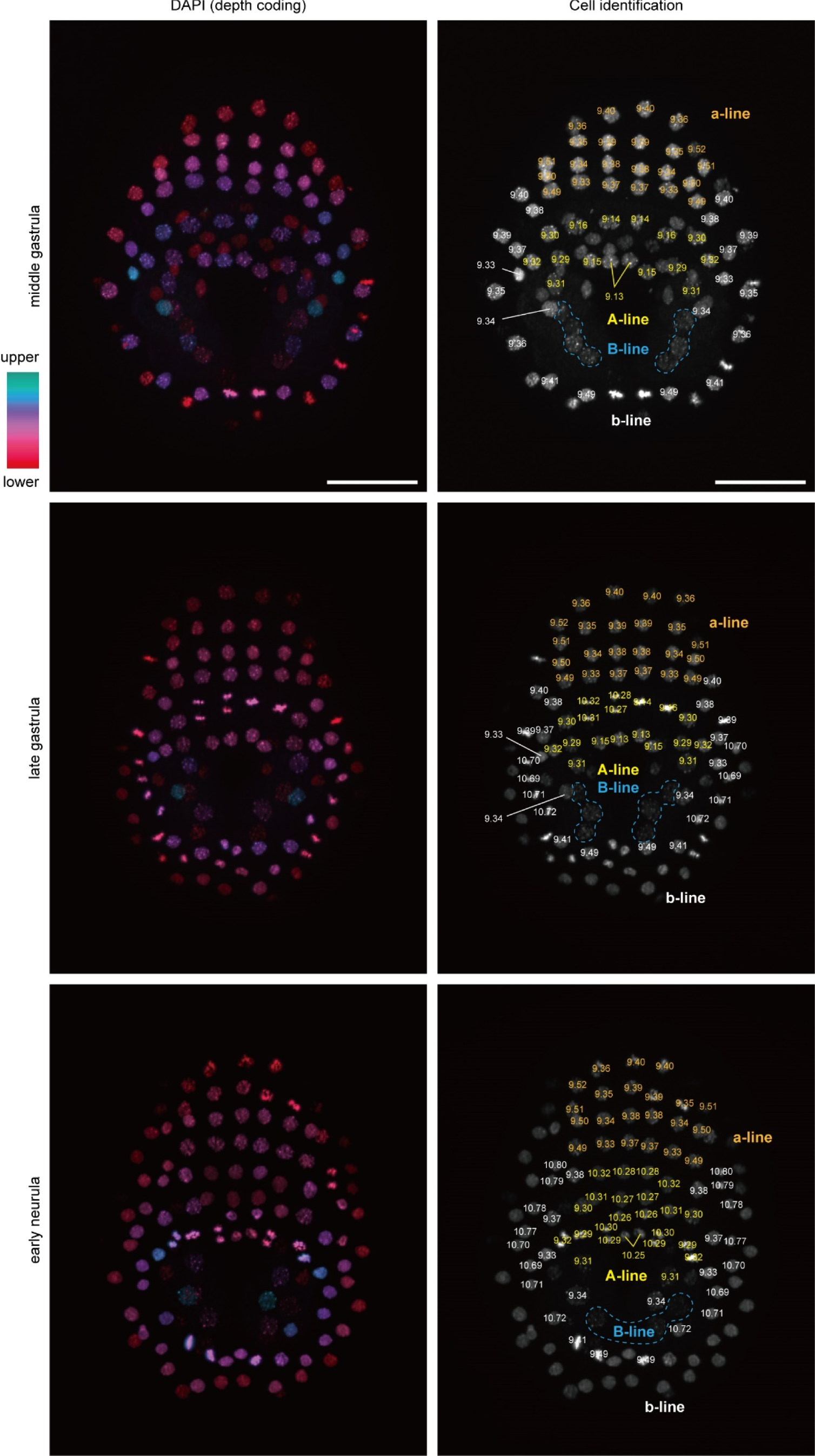
Cell identification from the middle gastrula to early neurula stages. Cell identities can be discriminated under the microscope. Nuclei were stained with DAPI, and photographs are z-projected image stacks overlaid. Color corresponds to the depth from the embryo surface in the left photographs. Cell names are shown on nuclei on the right photographs. Cells that are irrelevant to the present study are not necessarily identified.

